# Engraftment of wild-type alveolar type II epithelial cells in surfactant protein C deficient mice

**DOI:** 10.1101/2023.01.12.523571

**Authors:** Daniela Iezza, Camilla Predella, Keyue Ni, John W. Murray, Hsiao-Yun Liu, Anjali Saqi, Stephan W. Glasser, N. Valerio Dorrello

**Author notes:** Please address correspondence to, **N. Valerio Dorrello MD, PhD**, Associate Professor of Pediatrics, Division of Pediatric Critical Care and Hospital Medicine, NYP Morgan Stanley Children’s Hospital, Columbia University College of Physicians and Surgeons 3959 Broadway, CHN 10-13A (office);BB 421 (lab), New York, NY 10032, T.; F.

## Abstract

Childhood interstitial lung disease (chILD) secondary to pulmonary surfactant deficiency is a devastating chronic lung disease in children. Clinical presentation includes mild to severe respiratory failure and fibrosis. There is no specific treatment, except lung transplantation, which is hampered by a severe shortage of donor organs, especially for young patients. Repair of lungs with chILD represents a longstanding therapeutic challenge but cellular therapy is a promising strategy. As surfactant is produced by alveolar type II epithelial (ATII) cells, engraftment with normal or gene-corrected ATII cells might provide an avenue to cure. Here we used a chILD disease-like model, *Sftpc^−/−^* mice, to provide proof-of-principle for this approach. *Sftpc^−/−^* mice developed chronic interstitial lung disease with age and were hypersensitive to bleomycin. We could engraft wild-type ATII cells after low dose bleomycin conditioning. Transplanted ATII cells produced mature SPC and attenuated bleomycin-induced lung injury up to four months post-transplant. This study demonstrates that partial replacement of mutant ATII cells can promote lung repair in a mouse model of chILD.

## Introduction

Pulmonary surfactant lowers surface tension in the alveoli, thus preventing alveolar expiratory collapse and reducing the work of breathing. It is produced and recycled in lamellar bodies (LBs) of alveolar epithelial type II (ATII) cells starting at 24 weeks gestational age (1). Surfactant is composed of ~90% lipids of which 90% are phospholipids, and 8-10% surfactant proteins (SP) A, B, C, and D (2). 50–85% of surfactant is recycled by ATII cells while the remainder is degraded by alveolar macrophages (2–5). Children with mutations disrupting normal pulmonary surfactant production develop childhood interstitial lung disease (chILD) (6–11). ChILD secondary to surfactant defects is caused by mutations in *(i) SFTPB* (MIM: 178640), and *SFTPC* (MIM: 610913), which encode surfactant protein components; *(ii) ABCA3* (MIM: 610921), which encodes ATP-binding cassette sub-family A member 3, localized on the limiting membrane of LBs and essential for secretion; and *(iii) NKX2.1* (MIM: 610978), which encodes a transcription factor regulating *SFTPB*, *SFTPC* and *ABCA3* (6,7,9).

The incidence of ChILD ranges from 0.1 to 16.2 per 100,000 people (12,13) with a prevalence of 1.3-3.8 per million and mortality as high as 35% (6,9–11,14–16). Some forms of chILD are lethal in the neonatal period while others cause respiratory disease ranging from neonatal respiratory failure to childhood- or adult-onset interstitial lung disease. The pathology of chILD includes ATII hyperplasia, interstitial thickening, foamy macrophages, alveolar proteinosis, and interstitial pneumonitis (7,10). This pathology is explained at least in part by the fact that defective ATII cells affect alveolar homeostasis and cause fibrotic lung remodeling, as ATII cells are facultative progenitors of the alveolar epithelium after injury (17–24). In mice, for example, partial depletion of ATII cells increased susceptibility to bleomycin-induced fibrosis while total depletion of ATII cells caused extensive spontaneous lung fibrosis (25–27).

Supportive therapy, such as corticosteroids, hydroxychloroquine, and azithromycin has limited benefit for the majority of cases leaving lung transplantation as the *only* definitive treatment (6,9,28). Lung transplantation remains limited by the shortage of suitable donor organs, especially for smaller children (29,30). Moreover, the overall survival rate at five years post-transplant is just 55%, a figure that has not significantly improved in the last 20 years (30). There is therefore an urgent need for improved treatments. One study has demonstrated successful *in utero* gene editing using a mouse model of the most common *SFPTC* mutation responsible for chILD, *Sftpc^I73T^* (31). To be clinically applicable, diagnosis and correction have to occur before birth, which is not yet feasible. Furthermore, *SFTPC^I73T^* causes aberrant splicing, such that CRISPR-mediated splice site disruption is sufficient. Most surfactant deficiencies however are caused by missense and nonsense mutations, which are much more difficult to correct *in vivo*.

Cell therapy, in which gene-corrected cells are transplanted into the lungs to replace defective ATII cells, is an exciting treatment option for chILD. In rats, intratracheal (i.t.) administration of ATII cells after bleomycin injury may result in reduction of fibrosis (32,33). In mice infected with H1N1 influenza, oxygen saturation improved after ATII cell transplant (34). It is unknown whether transplanted ATII cells could affect the disease, and if so, how many endogenous cells need to be replaced with exogenous ATII cells for therapeutic effect. Eliminating all ATII cells may be neither safe nor feasible and is likely not necessary, as in many human diseases (e.g., hepatic, hematopoietic, and coagulopathic), not all the injured cells or enzymes need to be substituted to recover adequate function (35–38). The correct balance is unknown for ATII cell disease.

We investigated the efficacy of the cellular therapy in a viable mouse model of surfactant deficiency disease, the *Sftpc^−/−^* mouse. In humans, *SFTPC* encodes a 197 amino acids protein that is palmitoylated, proteolytically processed in the endoplasmic reticulum, and shuttled to the multivesicular bodies where final cleavage generating mature SPC (35 amino acids) takes place. SPC is then incorporated into LBs in which surfactant is stored prior to secretion in the alveolar space (2,39). *SFTPC* mutations can occur as missense, frameshift, insertion, deletion, and splice site mutations,and up to half are *de novo* mutations (40). The most common human *SFTPC* mutation is the threonine-to-isoleucine substitution at the codon 73 (*SFTPC^I73T^*) resulting in mistrafficking of SPC with proteotoxicity (41). In another subset of patients, no mutations in the *SFTPC* gene were identified, but SPC protein was still undetectable (42–44), suggesting that lung disease resulted directly from the absence of SPC protein. Clinically, mutations in *SFTPC* have variable penetrance with phenotypes ranging from neonatal respiratory failure to chILD to idiopathic pulmonary fibrosis in adulthood. Some patients require lung transplant, others are managed with long-term mechanical ventilation, and others remain almost asymptomatic. Postnatal mortality is rare (45,46).

In this proof-of-concept study, we show that cellular therapy is feasible with minimal lung conditioning in *Sftpc^−/−^* mice and that exogenous ATII cells can attenuate bleomycin-induced injury.

## Results

### Morphological changes of the lung in Sftpc^−/−^ mice

Previous work showed that *Sftpc^−/−^* mice on a 129/Sv background developed an interstitial pneumonitis-like phenotype beginning at two months of age and worsening with age up to 12-14 months (47). The phenotype was characterized by irregular alveolar septation, mononucleated cell infiltrates, ATII cell hyperplasia, and interstitial thickening (47). To develop a baseline time course of the disease, we analyzed the lung histology and stereology in *Sftpc^−/−^* mice at 4, 8, and 12 months of age **(Figure 1)**. Blinded histological analysis of representative lung sections **(Supplementary Figure 1)** showed progressive lung injury characterized by neutrophil infiltration into interstitium and airspace, alveolar wall thickening, and appearance of proteinaceous debris in the alveolar space. These features worsened with age in *Sftpc^−/−^* mice but not in *Sftpc^+/+^* controls **(Figure 1A)**. Lung injury scores (LIS), adapted from our work and others (48,49) and consistent with the American Thoracic Society (ATS) criteria for lung injury (50,51) were significantly higher (worse) in *Sftpc^−/−^* mice compared to age-matched *Sftpc^+/+^* mice **(Figure 1B)**. Alveolar septa count (number of interalveolar septa), volume density of alveolar septa (an estimation of alveolar airspace), mean trans-sectional wall length (a measure of alveolar septal thickness), mean linear intercept (estimation of volume to surface ratio of the airspace), and airspace surface area density (estimation of lung volume) were significantly altered in *Sftpc^−/−^* mice compared to *Sftpc^+/+^* mice at all ages **(Figure 1B)**, supporting the presence of a fibrotic remodeling process in *Sftpc^−/−^* mice consistent with prior data (47,52). The presence of interstitial thickening was further supported by elevated expression of vimentin, deposition of collagen (col) I and col IV, and increased hydroxyproline [(2S,4R)-4-hydroxyproline, Hyp] content in *Sftpc^−/−^* compared to *Sftpc^+/+^* lungs (**Figure 1C, Supplemental Figure 2A** and **B**). Within the *Sftpc^−/−^* mice group, the same parameters all significantly increased with age, but not in *Sftpc^+/+^* mice **(Figure 1C, Supplemental Figure 2A** and **B**). However, the total number of ATII cells did not appear to be affected in *Sftpc^−/−^* mice as immunostaining for Lysosomal Associated Membrane Protein 3 (LAMP3), an ATII cell marker, and expression of *Nkx2.1* and *Abca3* mRNAs, mainly expressed in ATII cells, were similar in *Sftpc^−/−^* and *Sftpc^+/+^* lungs **(Figure 1D)**. Immunostaining for SPC and expression of *Sftpc* mRNA were used as positive controls, since both expressed only in *Sftpc^+/+^* mouse lungs **(Figure 1D)**. Taken together, our data indicate that lack of SPC results in morphological and stereological changes of the *Sftpc^−/−^* mouse lungs similar to histological patters observed in chILD and worsening with age.

**Figure 1:**
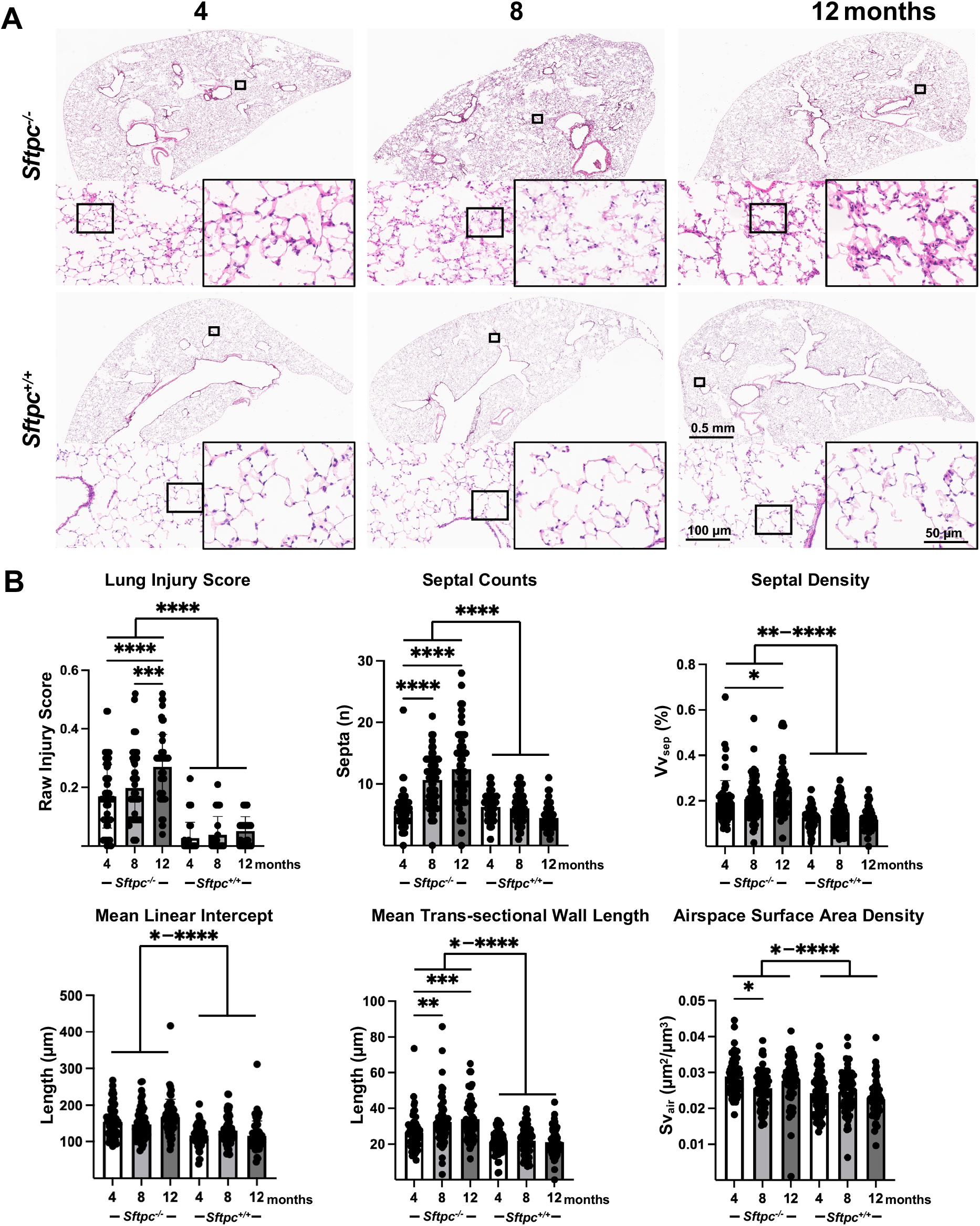
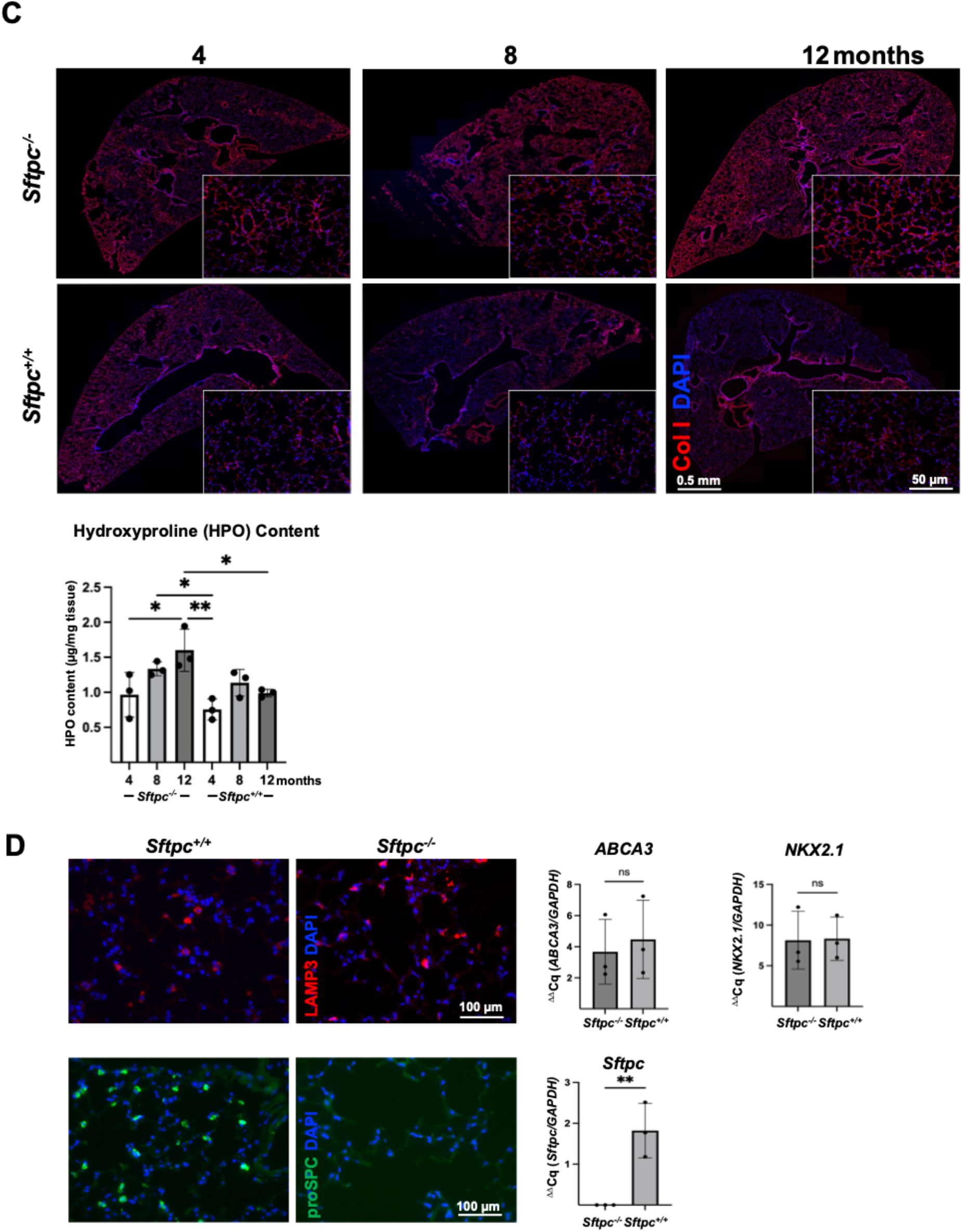
Morphological and stereological lung analysis of *Sftpc^−/−^* mice at different ages. **(A)** Representative histologic sections of whole lung comparing *Sftpc^−/−^* and *Sftpc^+/+^* mice at 4, 8, 12 months. Black squares indicate the area shown in the higher magnification. **(B)** LIS, morphometric, and stereological lung analysis are evaluated on 1000 × 1000 pixels lung histology fields (n=60 fields per group, 3 mice/group). LIS (***, p=0.0001; ****, p<0.0001). Septal counts (****, p<0.0001). Septal density (*, p=0.0104; **, p=0.0014; ****, p<0.0001). Mean trans-sectional wall length (*, p=0.0264; ***, p=0.004; ****, p<0.0001). Mean linear intercept (*, p=0.0394-0.0416; **, p=0.0073; ***, p=0.003; ****, p<0.0001) and airspace surface area density (*, p=0.0130-0.305; ***, p=0.0001-0.0004; ****, p<0.0001). Data are represented as mean ± SD and analyzed by one-way ANOVA. **(C)** collagen immunostatinig of representative lung sections of 4-,8-, 12-month-old *Sftpc^−/−^* and control mice (higher magnification in the boxed area; n=/group). Hydroxyproline content quantification via colorimetric hydroxyproline (HPO assay (y axis:HP content in μg/mg of lung tissue homogenates; x: ages for each group; n=3 per group, mean ± SD, one-way ANOVA; *, p=0.047-0.027; ** p=0035). **(D)** Expression by RT-qPCR of mRNA for alveolar epithelial markers *Lamp3, Abca3*, and *Nkx2.1*. Data are represented as mean ± SD and analyzed by unpaired two-tailed Student's t test (n.s., non-significant p>0.05; **, p≤0.0091). Data analyze *Sftpc* expression is absent in *Sftpc^−/−^* mice.

### Increased susceptibility to bleomycin in Sftpc^−/−^ mice

We next explored whether bleomycin could be used to ablate endogenous ATII cells. *Sftpc^−/−^* mice on a Black Swiss background have normal lung structure at baseline (53), but develop fibrosis after a lower dose (0.01U/mouse) of bleomycin compared to wild-type mice of the same background (54). In contrast, *Sftpc^−/−^* mice on 129Sv background, described here, show interstitial pneumonitis at baseline **(Figure 1** and (47))and develop extensive disruption of lung architecture and persistent lung inflammation after 0.05 U of bleomycin/mouse (a dose commonly used in wild-type mice) (55). To establish a dose response, we administered i.t. incremental doses of bleomycin (0.005, 0.01, and 0.05 U/mouse) i.t. to 4-month-old *Sftpc^−/−^* and *Sftpc^+/+^* mice **(Figure 2A)**. After 10 days, *Sftpc^−/−^* mice showed increased alveolar wall thickening, interstitial neutrophils, proteinaceous debris lining alveolar walls and filling airspaces at any dose of bleomycin while *Sftpc^+/+^* mice developed visible histological evidence of lung injury only at 0.05 U of bleomycin **(Figure 2B)**. Compared to *Sftpc^+/+^* mice, *Sftpc^−/−^* mice had significantly higher LIS at any dose **(Figure 2C)**. Stereological and morphometric analysis confirmed a bleomycin dose-dependent injury in *Sftpc^−/−^* mice, with increases in septal count, alveolar septal density, trans-sectional wall length, and mean linear intercept **(Figure 2C)**. While a similar trend was noticed in the *Sftpc^+/+^* mice, these changes were significantly less pronounced **(Figure 2C)**. Airspace surface area density was lower in *Sftpc^−/−^* mice regardless of bleomycin dose **(Figure 2C)**. Elevated deposition of collagens was observed only at the highest dose of bleomycin in the *Sftpc^−/−^* mice **(Figure 2D** and **Supplemental Figure 3A)**. Vimentin expression increased dose-dependently in *Sftpc^−/−^* mice but not in the *Sftpc^+/+^* mice **(Supplemental Figure 3B)**. Immunostaining for NKX2.1, used as a surrogate marker for ATII cells, showed a dose-dependent decrease of ATII in both *Sftpc^−/−^* and *Sftpc^+/+^* mice lungs **(Figure 2E** and **Supplemental Figure 4)**. RT-qPCR analysis confirmed the immunostaining data, showing downregulation of mRNAs encoding *Nkx2.1* as well as other ATII cell specific markers such as *Abca3* and *Sftpc* (expressed only in *Sftpc^+/+^* mice lungs) with increasing bleomycin dose **(Figure 2F)**. In the *Sftpc^−/−^* mice, *Nkx2.1* and *Abca3* mRNA levels were significantly low already at the lowest bleomycin dose (0.005U/mouse) **(Figure 2F)**. We conclude that *Sftpc^−/−^* mice are more susceptible to bleomycin injury than *Sftpc^+/+^* and that bleomycin depletes ATII cells.

**Figure 2:**
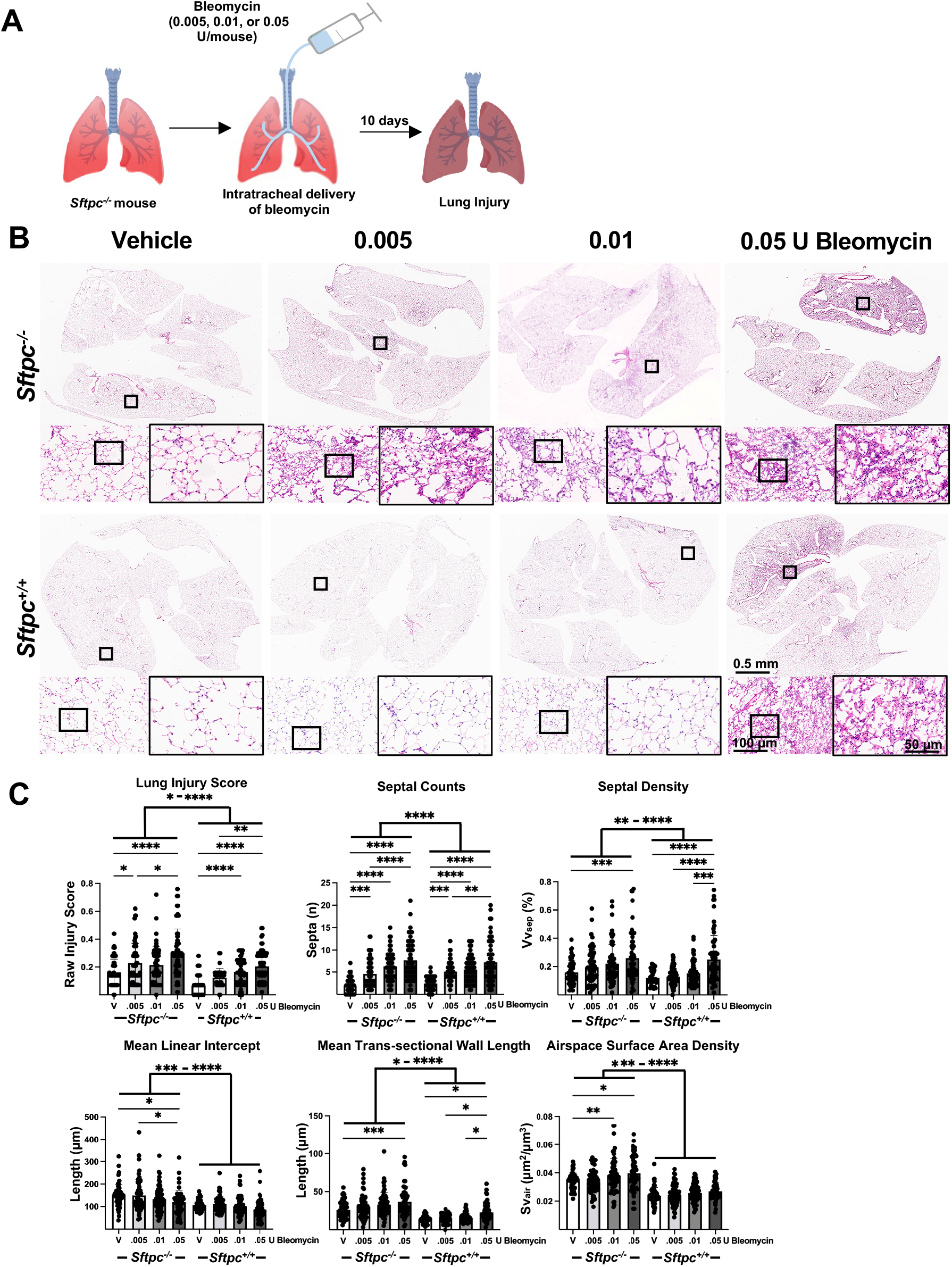
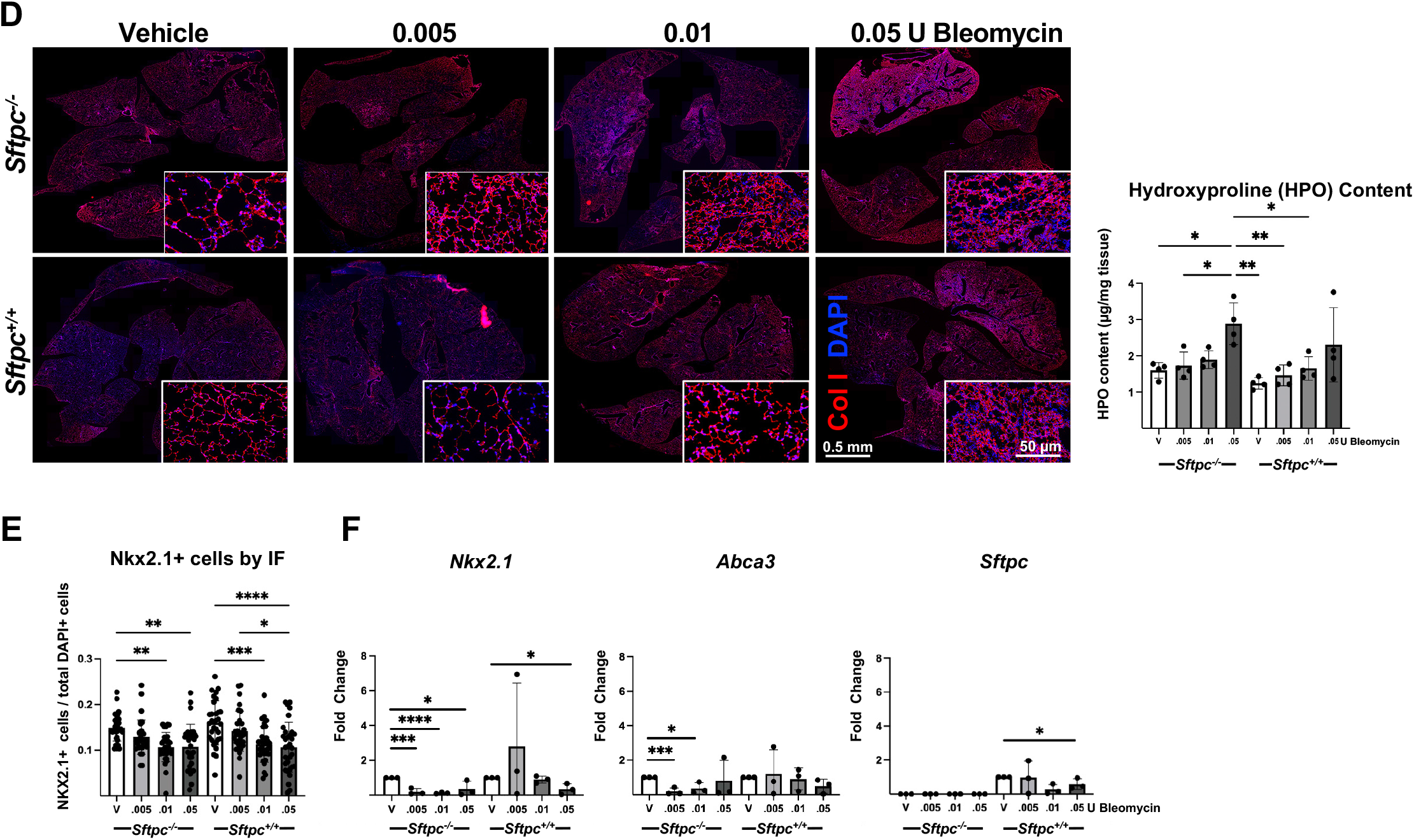
Susceptibility of *Sftpc^−/−^* mice to bleomycin. **(A)** Schematics of the injury model experiment with bleomycin. **(B)** Representative histologic sections of whole lung comparing 4-months-old *Sftpc^−/−^* and *Sftpc^+/+^* mice treated with increasing dose of bleomycin (0.005-0.05 U). Black squares indicate the area shown in the higher magnification. **(C)** LIS (*, p=0.0172-0.047; **, p=0.0039; ***, p=0.0002-0.0005; ****, p<0.0001), septal counts (*, p=0.0113; **, p=0.0051-0.0075; ***, p=0.0001-0.0005; ****, p<0.0001), septal density (*, p=0.0395; **, p=0.0015-0.0017; ***, p=0.0003-0.0009; ****, p<0.0001), mean trans-sectional wall length (*, p=0.0274-0.306; **, p=0.0095; ***, p=0.0005-0.0008; ****, p<0.0001), mean linear intercept (*, p=0.0125-0.0279; ***, p=0.0003; ****, p<0.0001) and airspace surface area density (***, p=0.0003-0.0005; ****, p<0.0001). All quantitative data are represented as Mean ± SD and statistically evaluated through one-way ANOVA. **(D)** Collagen I immunostaining of representative lung sections of 4-months-old of *Sftpc^−/−^* mice and control mice treated with 0.005, 0.01, or 0.05 U/mouse of bleomycin (higher magnification in the box area; n=3/group). Hydroxyproline content quantification via colorimetric hydroxyproline assay (n=3 per group; mean±SD, one-way ANOVA; *, p=0.0169-0.0417; **, p=0.0013-0.0066). **(E)** Nkx2.1 immunostaining quantification (count of Nkx2.1+ cells in random region of interests (ROI)s (n=20 per mouse, 3 mice/group. *, p=0.0299-0.0162; **, p=0.0025-0.0017; ****, p<0.0001). Data presented as mean±SD and statistically evaluated through one-way ANOVA with Tukey’s test. (**F)** Relative mRNA expression analyzed by RT-qPCR of *Nkx2.1, Abca3*, and *Sftpc* in lung homogenates of mice treated with different doses of bleomycin (*, p=0.036-0.0238 and ***, p=0.0026-0.0001 for *Nkx2.1*; *, p=0.018; ***, p=0.0001=0.0042 for ABCA3; *, p=0.0363 for *Sftpc*; n=3 per group. Data is represented by mean±SD and analyzed by one-way ANOVA.

### Engraftment of syngeneic Sftpc^+/+^ ATII cells in bleomycin-conditioned Sftpc^−/−^ mice

We next investigated engraftment of ATII cells in bleomycin-treated *Sftpc^−/−^* mice. Primary ATII cells were isolated from *Sftpc^+/+^* lungs, purified, and verified for epithelial markers and viability by fluorescence-activated cell sorting (FACS) (95-98% EPCAM^+^, CD45^−^) and immunostaining for pro-SPC **(Supplemental Figure 5A)**. 4-, 8-, and 12-month-old *Sftpc^−/−^* mice were conditioned with low-dose (0.005U/mouse) or high-dose (0.05U/mouse) bleomycin and 10 days later received 1×10^6^ *Sftpc^+/+^* ATII cells i.t. **(Figure 3A)**. In preliminary experiments, 1×10^6^ cells guaranteed consistent engraftment, 5×10^5^ cells resulted in fewer engrafted cells, while 2×10^6^ cells caused clumps of cells in the airways or inconsistent engraftment (data not shown). 14 days post-transplantation, lungs were harvested and analyzed for presence and extent of cell engraftment **(Figure 3A)**. By counting pro-SPC+ cells from immunofluorescent (IF) staining of representative lung sections of treated lungs, we noticed a higher engraftment of *Sftpc^+/+^* ATII cells in 4- and 8-month-old compared to 12-month-old mice **(Figure 3B and 3C)**. Better engraftment was observed at low dose bleomycin when compared to vehicle or to high dose **(Figure 3C)**. Pro-SPC counterstained with RAGE (ATI cell marker) showed ATII cells in proximity of ATI cells in their physiologic and anatomic context **(Figure 3B**, lower panel).

**Figure 3:**
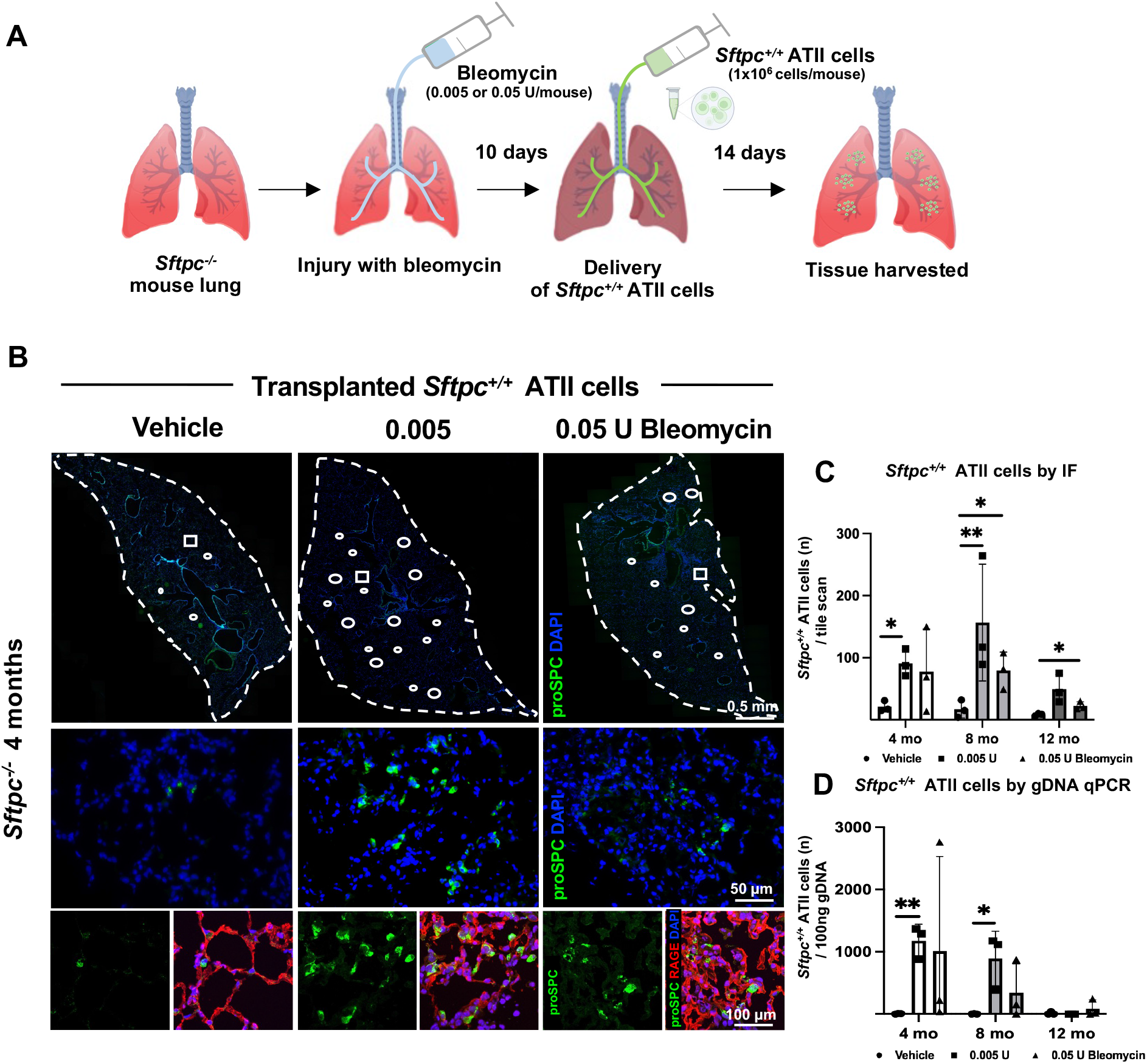
*Sftpc^+/+^* ATII cells engraftment in bleomycin-conditioned*Sftpc^−/−^* mice lungs. **(A)** Schematics of *Sftpc^+/+^* ATII cell transplant in *Sftpc^−/−^* mice post-bleomycin conditioning. **(B)** Immunostaining of lung sections for pro-SPC marker (ATII cell) 14 days post i.t. delivery of 1×10^6^ *Sftpc^+/+^* ATII cells in mice pre-conditioned with bleomycin (or vehicle) for 10 days. Lower panels: confocal images of alveolar regions showing the punctuated intracellular pattern of proSPC immunostaining within transplanted ATII cells surrounded by ATI cells, identified by the marker RAGE. Dotted line highlights the area of the analyzed lobes, round circles show areas where transplanted cells were found. White squares indicate areas shown in the higher magnification. **(C)** Pro-SPC immunostaining quantification (count of pro-SPC+ cells in ROIs (n=20 per mouse, n=3 per group. Data is represented by mean±SD, and analyzed by student t-test; *, p=0.031-0.036). Cell counts were as follows: 4-month-old (mo) mice (low dose bleomycin, 91±22cells/tile scan; high dose, 78±68 cells/tile scan; vehicle, 21±9 cells/tile scan. 8 mo mice (low dose bleomycin, 141±65 cells/tile scan; high dose, 80±30 cells/tile scan; vehicle, 13±8 cells/tile scan). 12 mo mice (low dose bleomycin, 56±35 cells/tile scan; high dose, 23±6 cells/tile scan; vehicle, 8±12 cells cells/tile scan). **(D)** Detection of *Sftpc^+/+^* cells in *Sftpc^−/−^* lung tissue via RT-qPCR for the *Sftpc* gene in 100 ng gDNA (3 sections per mouse; n=3 per group. Data is represented by mean ± SD, and analyzed by student t-test; *, p=0.0115-0.0186). Cells were detected as follows: 4 mo mice [low dose bleomycin, 1,181± 261 cells/100ng of gDNA; high dose, 1010±1520 cells/100ng of gDNA; vehicle, 7±6 cells cells/100ng of gDNA. 8 mo mice (low dose, 894± 439 cells/100ng of gDNA; high dose, 345±463 cells/100ng of gDNA; vehicle, 3±5 cells cells/100ng of gDNA). 12 mo mice (low dose bleomycin,17±28 cells/100ng of gDNA; high dose, 89±139 cells/100ng of gDNA; vehicle 14±4 cells/100ng of gDNA).

To confirm the pro-SPC immunostaining data we estimated the number of *Sftpc^+/+^* cells transplanted in the *Sftpc^−/−^* lungs by RT-qPCR for *Sftpc* gene. 100 ng of genomic (g) DNA was extracted from 200μm thick lung sections derived from transplanted mice and subject to RT-qPCR analysis. Using a methodology previously applied for other transplantation models (56), we compared the *Sftpc* qPCR values of the our samples to a standard curve obtained by the qPCR values derived by known number of *Sftpc^+/+^* cells **(Supplemental Figure 4B)**. Again, we detected the higher number of transplanted *Sftpc^+/+^* cells in younger mice (4 and 8 months) compared to older mice (12 months) and after low compared to high bleomycin dose or vehicle **(Fig. 3D)**. Overall, low-dose of bleomycin, while causing a minimal lung injury, is permissive for a robust *Sftpc^+/+^* ATII cell engraftment especially in younger *Sftpc^−/−^* mice, while higher dose might cause severe lung injury that impairs efficient and consistent engraftment at any age.

### Long-term engraftment and functionality of Sftpc^+/+^ ATII cells

Next, we asked if primary *Sftpc^+/+^* ATII cells could engraft for prolonged periods and promote repair. 4-month-old *Sftpc^−/−^* mice were conditioned with low-dose bleomycin and 1×10^6^ *Sftpc^+/+^* ATII cells were delivered i.t. 10 days later, lungs were harvested and analyzed 2, 4, 8, and 16 weeks post-transplantation. We could detect a significant amount of pro-SPC+ ATII cells in the alveolar region of all the *Sftpc^−/−^* mice conditioned with bleomycin compared to those conditioned with vehicle only prior to transplant **(Figure 4A** and **4B)**. By RT-qPCR for *Sftpc*, we confirmed the persistence of *Sftpc^+/+^* cells up to 16 weeks **(Figure 4C)**. These data suggested that conditioning with bleomycin allowed both short- and long-term engraftment **(Figure 4A-C)**. From 2 weeks on, we could detect patches of LAMP3+ SPC+ ATII cells, demonstrating that transplanted *Sftpc^+/+^* ATII cells were viable and capable to fully process pro-SPC in its mature product, SPC, in the recipient lungs **(Figure 4D)**.

**Figure 4:**
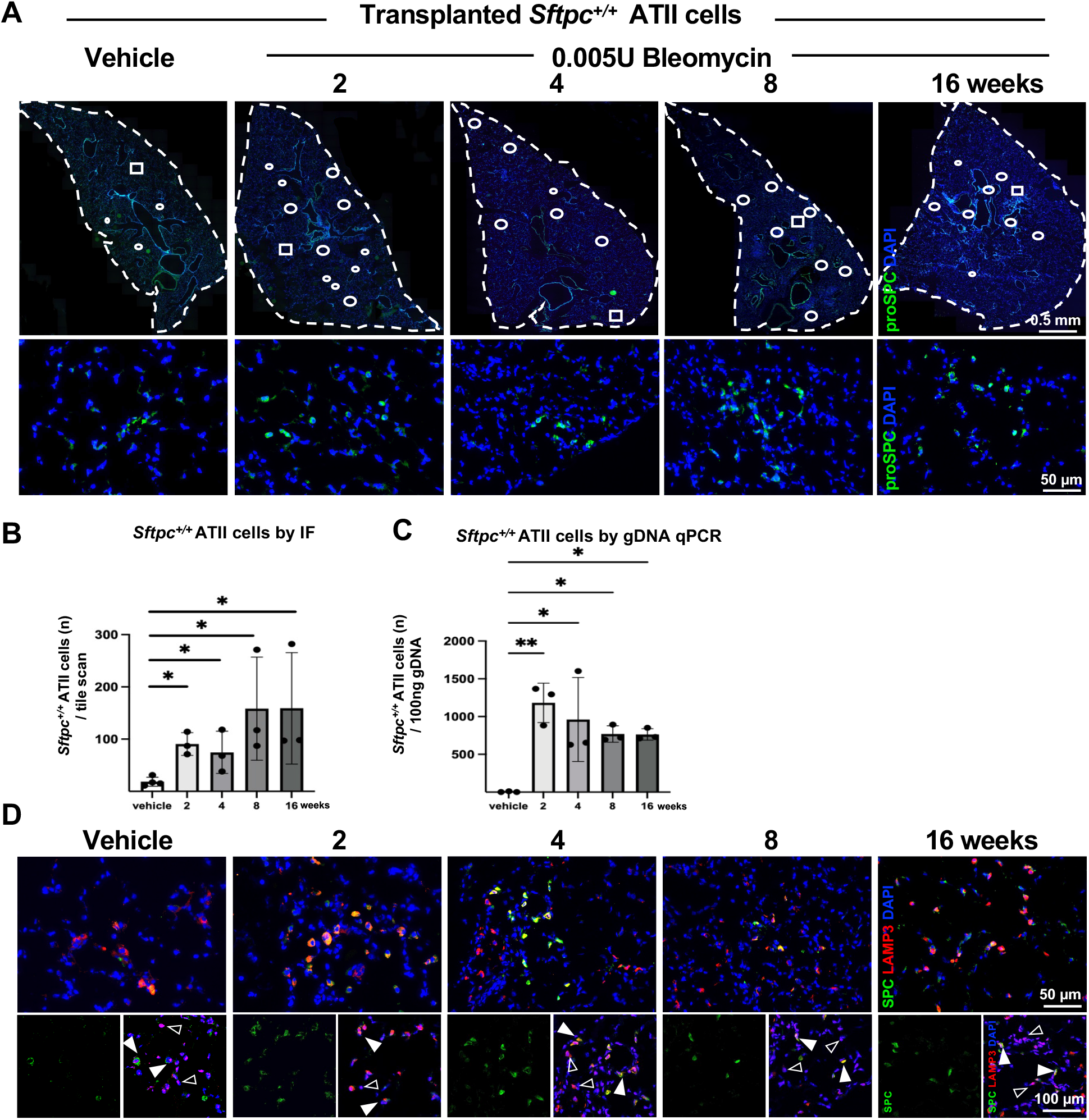
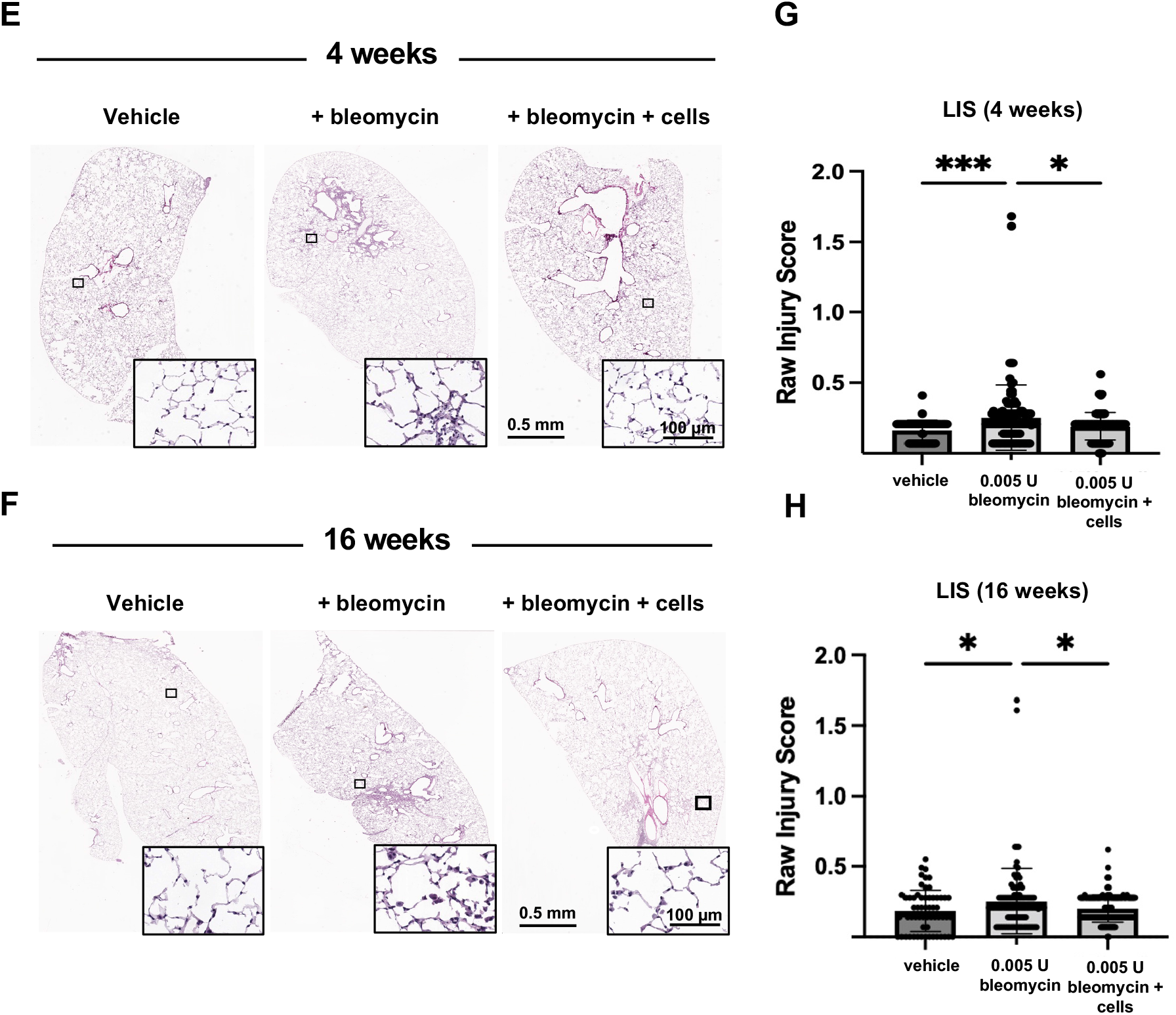
Long-term viability and functionality of transplanted *Sftpc^+/+^* ATII cells in *Sftpc^−/−^* mice. **(A)** Representative images of whole lung tile scans immunostained for pro-SPC 2, 4, 8, and 16 weeks post-transplantation of *Sftpc^+/+^* ATII cells in mice pre-conditioned with bleomycin (or vehicle) for 10 days. Dotted while line highlights the area of the analyzed lobes, round circles shown areas where transplanted cells were found. The white squares indicate the area shown at higher magnification. **(B)** Pro-SPC immunostaining quantification (count of pro-SPC+ cells in whole lung tile scans (n=3 per mouse, n=3 per group). Data is represented by mean ± SD and analyzed by one-way ANOVA; *p=0.027). Counted cells are as follows: 14±3, 91±22, 75±40, 158±99, 159± 107 cells/tile scan for vehicle, 2, 4, 8, and 16 weeks, respectively. **(C)** Detection of *Sftpc^+/+^* cells in *Sftpc^−/−^* lung tissue via RT-qPCR for *Sftpc gene* in gDNA (3 sections per mouse; n=3 per group). Estimated cell numbers are as follows: 4±6, 1182±261, 961±556, 769±109, 763± 76 for vehicle, 2, 4, 8, and 16 weeks, respectively. Data is represented by mean ± SD and analyzed by one=way ANOVA; **, p=0.0016-0.0066). **(D)** mature SPC counterstained with LAMP3 (ATII cells) showing engrafted *Sftpc^+/+^* ATII cells (LAMP3+ SPC+) are capable of producing mature SPC in the *Sftpc^−/−^* lungs. Representative LAMP3+ SPC+ cells are indicated by full arrows. Representative LAMP3+ SPC-cells (*Sftpc^−/−^* ATII cells) are indicated by empty arrows. **(E and F)** Representative histological images of whole lung sections at 4 (E) and 16 (F) weeks of transplanted *Sftpc^−/−^* mice and time-matched controls (vehicle or bleomycin and mock cell delivery). **(G and H)** LIS is calculated for 4 weeks (G) and 16 weeks (H). (60 lung histology fields per group, 3 mice/group. Data is shown as Mean ± SD and analyzed by one-way ANOVA; *, p=0.0140-0.0218, ***, p=0.003).

Finally, we asked whether transplanted ATII cells could promote repair of lungs injured by bleomycin. Histological analysis of mice lungs at 4 weeks and 16 weeks post-transplantation suggested improved lung structure and morphology when compared to mice conditioned with bleomycin but with mock cell delivery **(Figure 4E** and **F)**. At both time points, transplanted mice had an improved LIS compared to mice treated only with bleomycin, and similar to vehicle-treated mice **(Figure 4G** and **H)**. Taken together, these data indicate that transplanted ATII cells can attenuate lung injury induced by bleomycin.

## Discussion

Using a chILD-like disease model, the 129/Sv *Sftpc^−/−^* mouse, *Sftpc^+/+^* ATII cells can engraft after pre-conditioning with low-dose bleomycin, especially early in the course of the disease, and engrafted *Sftpc^+/+^* ATII cells are capable of repairing lung injury secondary to bleomycin.

The initial study of Glasser and colleagues reported the generation of SPC null mutant mice in a Swiss black background (53). Those mice, despite lacking SPC, were viable at birth and developed normally with only mild alterations in lung mechanics and decrease stability of surfactant when analyzed *in vitro*. In contrast, the same mutation on a 129/Sv background, used for the current studies, induced ATII hyperplasia with LB-like and lipid inclusions, increased neutrophils, alveolar macrophages with intracellular surfactant-like material, accumulation of α-smooth muscle actin (α-SMA) and are, therefore, a better model of the human disease (47). Importantly, these mice normal at birth, subsequently exhibit signs of interstitial fibrosis and cellular inflammation starting at 2 months of age and progress with age with extensive remodeling, airspace loss and patchy fibrosis (47). Similarly, *Sftpc^−/−^* mice were more susceptible to respiratory syncytial virus (RSV) infection as well as to bacteria, with delayed resolution of the pulmonary alterations (57–59). Other animal models carrying *SFTPC* mutations exist, including a mouse expressing most common *SFTPC* mutation found in human patients, *SFTPC^I73T^* (17). The *Sftpc^I73T^* mouse model*^,^* results in embryonic lethality unless inducibly expressed in the post-natal period (17,60). After tamoxifen induction, these mice develop severe polycellular alveolitis with increased mortality between 7 and 14 days. This is in contrast with children affected by chILD-*SFTPC^I73T^* in which symptoms appears variably after birth and not typically associated with postnatal mortality (61,62). Other variants of *SFTPC* in humans include L188Q, Δexon4, and C121G, all localized to the pro-SPC COOH-terminal (BRICHOS) domain (18,63). They cause protein misfolding, aggregation, ER stress and apoptosis (18,63). Mice expressing Δexon4 *SFTPC* transgene or the C121G mutant show ATII cell death, fibrotic remodeling, and neonatal lethality (64,65). When C121G mutant is expressed in an inducible manner, instead, mice show a pattern of chronic fibrosis (50). The transgenic mouse expressing L188Q and a recent knock-in model expressing this mutation, do not develop lung fibrosis unless conditioned with bleomycin (66,67). In contrast to these inducible models, the model used in the current study showed a consistent and well-defined phenotype across the experiments **(Figure 1)**.

Bleomycin remains the most frequently used agent to experimentally induce pulmonary fibrosis in animal models. Bleomycin injury however does not faithfully recapitulate the human fibrotic lung disease as it induces an inflammatory fibrosis that resolves over time (68). We used low-dose bleomycin with goal of depleting ATII cells prior to cell transplant. In our case, 0.005U/kg of bleomycin (one tenth of the dose normally used in wild-type mice) was sufficient to partially remove ATII cells without severely further damaging lungs **(Figure 2)**. While bleomycin fits the purpose of animal studies, and is commonly used as pre-conditioning strategy, we are aware that alternative alveolar methods need further investigation to be applicable to humans.

When considering cell therapy, the first question is the choice of engrafting cells. Recent studies have shown that both human and murine primary, embryonic or pluripotent stem-cell derived cells can be transplanted in recipient murine lungs post injury and cells persisted up to 4-6 months (34,69–75). In all these previous studies, wild-type or immunodeficient mice were used as recipient. We used highly purified *Sftpc^+/+^* ATII cells as a proof-of-concept study and a disease model as recipient. Primary ATII cells have been previously used in transplantation experiments in wild-type mice showing engraftment and certain degree of whole organ recovery post injury (34). The cells transplanted in our study present several advantages. They are primary ATII cells isolated from *Sftpc^+/+^* mice lungs and they express SPC. Therefore, when transplanted in *Sftpc^−/−^* mice, they can potentially correct the phenotype of the host, reintroducing SPC. They are easily identifiable in the *Sftpc^−/−^* mice by immunofluorescence for pro-SPC or SPC and by RT-qPCR for *Sftpc* facilitating any engraftment analysis without requiring external manipulations for identification (e.g. lentiviral fluorescent labeling). Finally, since they are syngeneic, the host does not require any immunosuppression to allow engraftment. We note that murine Pluripotent Stem Cell (mPSC)-derived cells could be used here. However, generating ATII cells from mPSCs has been very challenging, likely because of the very narrow developmental time windows in mouse development compared to human development. Multipotent Sox9^+^ lung progenitors have been isolated from fetal lungs, expanded in 3D culture, and showed to engraft in immunocompromised mice after injury and differentiate in airway and alveolar cells (76). Recently, another group has been able to derive and expand *in vitro* Nkx2-1+/Sox9+ lung epithelial progenitors from murine embryonic stem cell line and showed differentiation into ATII-like and ATI-like cells when transplanted into the lungs of syngeneic immunocompetent recipients (71). While these results are excited and promising, they have not yet been applied to a disease model nor evaluated for therapeutic effect on disease progression.

A second question is the determination of the number of defective cells to be replaced to demonstrate a therapeutic benefit for patients. Replacing all ATII cells may be neither safe nor needed. For example, in *Sftpb^−/−^* mouse models, symptoms appear only when SPB levels fall below 20-25% of wild-type levels (77). Our data support the idea that partial replacement of ATII cells is sufficient to promote lung repair, as we transplanted up to 1×10^6^ ATII cells, one tenth of total ATII cell number in a mouse lung (78), which was sufficient to attenuate lung injury secondary to bleomycin **(Figure 4E** and **F)**.

The third question is timing of intervention. In chILD as well as in the adult ILD, there is an urgent need for both early diagnostic and therapeutic strategies. All the current therapies aim at attenuating the downstream effects of the disease but do not target the main cell involved in its pathogenesis, the ATII cell (18,79). In replacing or correcting defective ATII cells, timing is critical since it should ideally occur prior to irreversible chronic changes of the lung. In this study, we transplanted animals at 3 different ages, (4, 8, and 12 months of age) corresponding to three stages of progressive lung disease and using bleomycin as pre-conditioning strategy. We noticed that at 4 and 8 months of age, low dose of bleomycin had the higher cell engraftment between all the conditions tested (**Figure 3)**. In 12-month-old mice, instead, cell engraftment was always lower for any bleomycin dose and variable between experiments. This could be explained by the fact that the natural progression of lung disease of the *Sftpc^−/−^* mice aggravated by bleomycin creates a hostile milieu for cell attachment and engraftment. Our data suggested that cell therapy for progressive lung diseases such as chILD should occur early in the disease course. In a vulnerable population such as children with chILD, we envision a regional multi-step approach, where pockets of lung get treated with cell therapy one by one until acceptable lung function is achieved.

We observed long-term engraftment and attenuation or repair of bleomycin-induced injury (**Figure 4D-F)**. Our data support those of other groups on mouse-to-mouse cell transplants using bleomycin as injury mechanism and i.t. administration as delivery route (34,72,73). Louie and colleagues showed that Rag1 KO mice conditioned with bleomycin and transplanted with organoid cells derived from SCA1-negative ATII cells had lower Ashcroft injury scores compared to those injured only with bleomycin (73). In another study, club-like lineage negative epithelial progenitors with high H2-K1 expression were transplanted in mice conditioned with bleomycin. While histology was not evaluated for lung repair, they did notice improve in oxygen saturation at 2 weeks post-transplant (72). In a similar study, mice conditioned with bleomycin and transplanted with primary ATII cells improved their oxygen saturation and their arterial partial pressure of oxygen at 2 weeks post-transplant (34). Regardless of the mechanisms and signals involved, the current and other studies (34,72,73) showed that transplanted ATII or ATII-like cells could promote repair or functional recovery of the lung. Future studies should investigate the transcriptomic and proteomic signatures dictated by ATII derived cells transplanted, as well the long-term fate of these cells.

Our study has several limitations. While bleomycin is commonly used as pre-conditioning strategies (34,71–73), this conditioning approach is not applicable to humans. Another limitation is that it is challenging to accurately determine the total number of engrafted *Sftpc^+/+^* ATII cells in the whole mouse lung. While *Sftpc^+/+^* ATII cells could be identified by pro-SPC immunostaining, the lack of lineage tracing of our transplanted cells did not allow us to follow the fate of these cells or determine if they differentiated into other lung cell types, including ATI cells. This could lead to underestimate the number of cells and types derived from the initially engrafted *Sftpc^+/+^* ATII cells.

Nonetheless, our study demonstrates the successful engraftment and persistence of ATII cells in a chILD-like disease model. Using low dose bleomycin as conditioning strategy, we showed that primary ATII cells could engraft and ameliorate bleomycin induced injury in those mice. Ultimately, *Sftpc^−/−^* mice with a minimal conditional strategy, such bleomycin, could represent a preclinical, translational platform to assess cell therapy in chILD. The potential use of human Pluripotent Stem Cell-derived lung progenitors as engrafting cells (80–82) in a chILD-like disease animal model as described here under adequate pharmacological immunosuppression could represent an attractive and promising avenue with rapid clinical translation. In conclusion, our study lays the foundation for cell therapy in chILD, offering an alternative approach to lung transplantation.

## Methods

### Animals

129S2/SvPasOrlRj (129/Sv) *Sftpc^−/−^* mice, generated by gene inactivation as previously described (47), were generously donated by Dr. Whitsett, Cincinnati Children’s Hospital. 129/Sv *Sftpc^+/+^* (wild-type) mice used as control group were purchased by Taconic Biosciences (NY, USA). In all the experiments, 4-, 8-, or 12-month-old mice, both female and male, were used. All animal work was approved by the Columbia University Institutional Animal Care and Use Committee and complied with the National Research Council *Guide for the Care and Use of Laboratory Animals*. Mice were humanely euthanized via inhalation of 5% isoflurane, followed by a second method of euthanasia: cervical dislocation followed by median sternotomy. Lungs were perfused via right ventricle with PBS (Phosphate Buffer Saline), and following perfusion, an incision was made, and the trachea was cannulated using a 20G catheter. The catheter was secured in position via a suture wire, and the lungs were filled by intratracheal instillation of 80% OCT (optimal cutting temperature compound, Sakura Finetek)/20% PBS, removed surgically from the animal, frozen, sectioned according to pre-determined map **(Supplementary Figure 1)**, and stored at -80°C.

### Delivery of Bleomycin and Induction of Lung Injury

15 Units (U) of bleomycin powder (Meitheal Pharmaceuticals Inc.) was resuspended in 5 mL of sterile normal saline (NS) solution and stored sterile at 4°C. For any use, only the necessary amount was taken from the vial. 4-12 months-old 129/Sv *Sftpc^+/+^* and *Sftpc^−/−^* 129/Sv mice were weighed and anesthetized with an intraperitoneal injection of Ketamine (80-95 mg/kg) and Xylazine (5-10 mg/kg). Intratracheal intubation was performed using a 20G cannula modified in length for murine lungs. Bleomycin was administered at 0.005, 0.01, or 0.05 U/mouse in a total volume of 40μL in sterile NS. Control mice received 40μL of vehicle (NS). Following i.t. administration, mice were allowed to recover from anesthesia and returned to their cages until euthanasia. Endpoint analysis was performed at 10 days from bleomycin injection. Mice were humanely euthanized with isoflurane and lung tissues were harvested, frozen in OCT, and stored at −80°C. Tissues were processed for Hematoxylin and Eosin (H&E), immunofluorescent staining, or gDNA/mRNA extraction.

### Stereological Analysis and Lung Injury Score

H&E sections were prepared by the Herbert Irving Cancer Center Molecular Pathology Core. Sections were scanned on a Leica AT2 slide scanner at 40X resolution (.25 microns/pixel). 1000 x 1000 μm sections were randomly generated using a ImageJ macro available (https://github.com/diezza1832/codes-for-Iezza-et-al-2022/blob/main/Random section generator.ijm) and then were analyzed for morphometric and stereological features, and LIS. For each analysis, 60-120 random sections were analyzed for each condition. Vvsep, Vsair, Lm, and Lmw were calculated according to the methods previously described (48,83). Septal counts were performed manually by two independent researchers blind to the treatment group using the grid system previously described by our group (49). For LIS, these same sections were analyzed by a pathologist blinded to the treatment group according to ATS guidelines (50,51) to determine lung injury. Further details regarding pathological analysis and criteria can be found in Supplementary materials **(Supplemental Table 1)**.

### Hydroxyproline Assay

Hydryoxproline content was determined for each sample using Millipore Sigma Hydroxyproline Assay kit (MAK008). 25-30 mg of tissue was sectioned from tissue-blocks cryopreserved in OCT. Sections were washed in PBS to remove residual OCT, weighed, dried, hydrolyzed, and ran according to manufacturer’s protocol.

### Isolation of ATII Cells

Healthy 129/Sv *Sftpc^+/+^* mice at 2-3 months of age were humanely euthanized for isolation of *Sftpc^+/+^* ATII cells. Lungs were perfused with PBS via the right ventricle of the heart and bronchioalveolar lavage was performed with PBS prior to inflate lungs *in situ* with Dispase (~0.9 ml, 50U/ml, Corning #354235) via tracheal cannulation. Lungs were tied with suture wire and incubated in an additional 1 mL of Dispase at room temperature (RT) for 20 min. Following incubation, pulmonary lobes were separated from the trachea and bronchial tree and chopped mechanically with a variety of surgical scissors. After dissection and mechanical separation, the tissue was incubated at 37°C in MEM + DNase I with frequent agitation to complete digestion for 10 min. The mixed cell digest was filtered through 100 μm and 40 μm cup filters, and fibroblasts were removed from the suspension by three successive adherence steps on tissue-culture treated plastic. Alveolar epithelial type II cells were then purified from resident macrophages, lymphocytes, and blood cells through two passes of cell enrichment and isolation kits (Dynabeads DC Cell Enrichment Kit #114.29D; Dynal Mouse T Cell Negative Isolation Kit #114.13D). Kits were used as described in manufacturer protocol. The purity of the resulting ATII population was verified both via immunostaining (pro-SPC) after plating overnight the cells in Matrigel (1:30 mixed in media), and by selecting CD45-EpCAM+ population (CD45 BV421, Biolegend #103133; EpCAM PE/Cy7, Biolegend #118215) using a Sony MA900 cell sorter (San Jose, CA).

### Bleomycin Conditioning and ATII Cell Engraftment

4-, 8-, or 12-months old *Sftpc^−/−^* 129/Sv mice were anesthetized and treated with 0.005-0.05 U of bleomycin, or vehicle in 40μL volume via i.t delivery as previously described in this paper. Each age group contained 3 animals for each treatment condition. After 10 days, mice received 1×10^6^ freshly isolated *Sftpc^+/+^* ATII cells delivered i.t. and were harvested at 2-16 weeks post cell delivery. Lung tissues were inflated with 80% OCT/20% PBS and sectioned into upper, middle, and lower lobes according to a predetermined lung map that covers both lungs from upper to lower regions **(Supplemental Figure 1)**.

### Immunofluorescent Staining

Lung sections were thaw at RT for 5-10 minutes, fixed with 4% paraformaldehyde for 10 minutes at room temperature (RT) and washed with PBS for 5 minutes. The sections were permeabilized with 0.25% Triton X-100/PBS for 20 minutes followed by blocking in 10% donkey serum diluted in PBS for 1 hour. Primary antibodies **(Supplementary Table 2)** were diluted in 5% donkey serum in PBS and incubated at 4°C overnight. The following day, sections were washed with PBS + 0.025% Triton-X for three times, 5 minutes each followed by secondary antibody **(Supplementary Table 3)** incubation for 1 hours at RT. Following secondary incubation, sections were washed for three times for 10 minutes with PBS + 0.025% Triton-X100 and finally mounted with DAPI contained fluorescent mounting medium. The following antibodies were used: proSPC (Seven Hills, #WRAB-9337), mature SPC (Seven Hills, #WRAB76694), RAGE/AGER (R&D, #AF1145), Collagen I (abcam, #ab34710), Collagen IV (abcam, #ab6586), vimentin (abcam, #ab92547), LAMP3 (Synaptic Systems, #391005), and anti-TTFI (Seven Hills, WRAB-1231), Donkey α Rabbit IgG 488 (Invitrogen, #A21206), Donkey α Rabbit IgG 555 (Invitrogen, #A31572), Donkey α Rabbit IgG 647 (Invitrogen, #A31573), Donkey α Goat IgG 568 (Invitrogen, #A11057), and Goat α Guinea Pig IgG 555 (Invitrogen, #A21435) with the dilutions according to the tables **(Supplementary Tables 2** and **3)**. For proSPC and mature SPC staining in the transplant experiments analysis,TrueVIEW® Autofluorescence Quenching Kit (Vector, SP-8400-15) was applied and counterstained with DAPI prior to imaging. Samples were imaged using motorized DMi8 (Leica Microsystems, Buffalo Grove, IL) inverted microscopes. Confocal images were taken on a Leica Stellaris 8. For NKX2.1 stainings, 20 random 500 x 500 μm regions of interest were generated for each sample and positive cells were counted over total number of cells.

### RT-qPCR

200 μm thick slices of lung tissue cryopreserved in OCT were sectioned and washed with PBS to remove residual OCT. For mRNA extraction, tissue was digest and homogenized in Trizol reagent according to manufacturer’s protocol (Invitrogen™, #15596026). For RT-qPCR of mRNA expression, mRNA was extracted using Zymogen Direct-zol RNA Microprep kit (#R2062), and cDNA was generated from RNA extracts using the Multiscribe High-Capacity cDNA kit (ThermoFisher, #4368814). For the analysis, 20 ng of cDNA was analyzed per well. For genomic DNA extraction, tissue was lysed, and gDNA was extracted using Zymogen Quick-DNA Microprep Kit (#D3021) according to manufacturer’s protocol. For the analysis, 100 ng gDNA was analyzed per well. ΔΔ Comparative qPCR was run on all samples using SYBR green reagents (ThermoFisher, #A25743) and using gene-specific PCR primers from Integrated DNA Technologies **(Supplementary Table 4)**.

## Data availability

The data that support the findings of this study are available on request from the corresponding author, NVD.

## Statistical methods

Statistical methods applied for each experiment are outlined in the figure legends. Both male and female mice were used with no clear difference in response to bleomycin or transplantation. Unpaired, two-tailed Student’s t tests were used when comparing two groups, while ANOVA and Tukey’s test were used when comparing multiple groups. For each data set, mean±SD were calculated and presented in the legend section of each figure. The differences between the groups were considered statistically significant for p ≤ 0.05.

## Study approval

Mice were housed in a pathogen-free barrier facility at Columbia University Vagelos College of Physicians and Surgeons. All animal procedures were performed under protocol (RASCAL AC-AABP05353) approved by the Institutional Animal Care and Use Committee of Columbia University Vagelos College of Physicians and Surgeons.

## Author contributions

DI, CP, and NVD designed the study. DI, CP, SG, HL, and NVD interpreted data. NVD wrote the manuscript with support of DI, CP, SG. DI, CP, KN conducted the experiments and analyzed data. AS performed the blinded pathologic assessment. JWS assisted with confocal imaging, H-YL assisted with gDNA RT-qPCR. All the authors read and approved the final manuscript.

## Acknowledgments

The authors thank the following collaborators and supporters: Hans-Willem Snoeck for fruitful discussion of the data and reading the manuscript; Joshua Matelow for critical reading of the manuscript; Institute of Comparative Medicine veterinary staff including Rivka L. Shoulson for supporting animal studies; Theresa Swayne at Confocal and Specialized Microscopy Shared Resource of the Herbert Irving for analytical support; the Herbert Irving Comprehensive Cancer Center Confocal and Specialized Microscopy Shared Resource, funded in part through NIH/NCI Cancer Center Support Grant P30CA013696; Michael Kissner and Columbia Stem Cell Initiative for assistance with flow and sorting analysis. The authors gratefully acknowledge funding support from the Department of Defense (PR180834, NVD), Driscoll Children’s Fund Scholar (NVD), and Louis V. Gerstner Jr. Scholarship Fund (NVD).

## Supplemental Materials

**Supplementary Figure 1:**
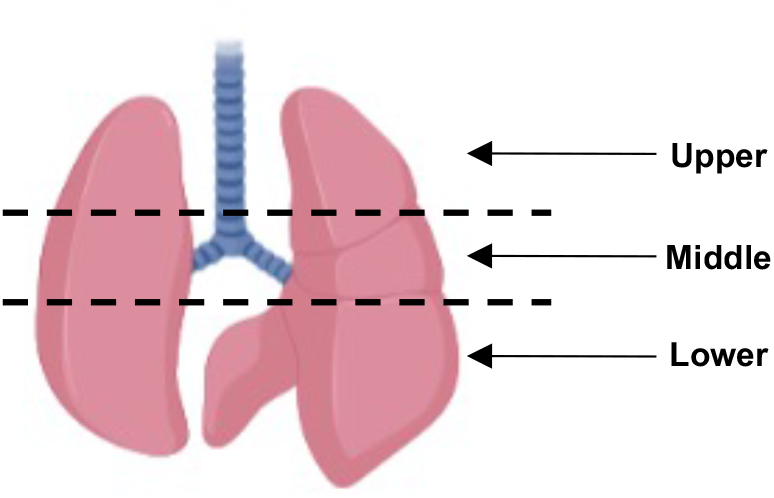
Lung map used for sampling. Lung was divided into 3 regions— upper, middle, and lower— each one including left and right sides of the lung.

**Supplementary Figure 2:**
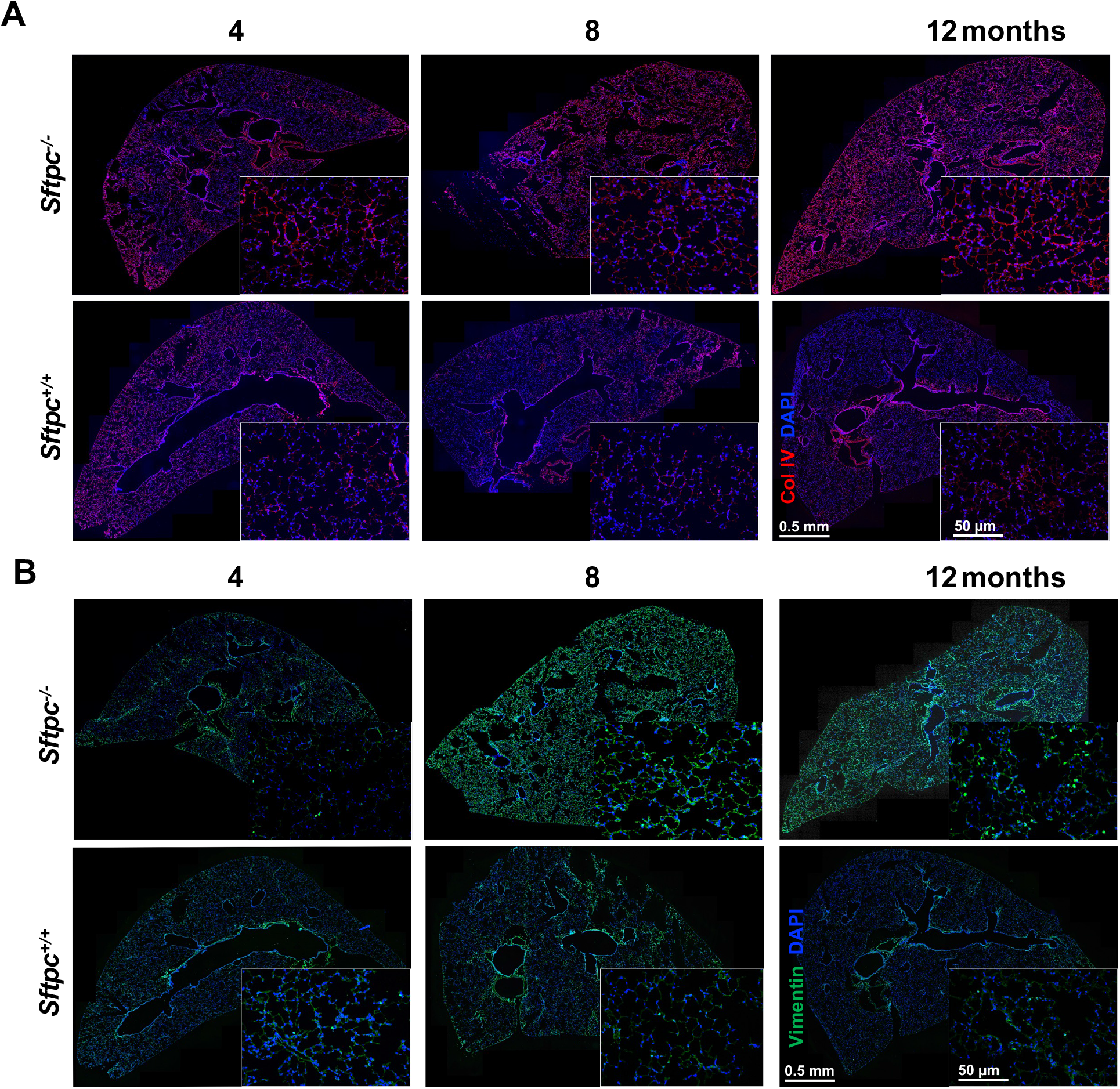
*Sftpc^−/−^* mice develop lung morphological changes with age. **A** and **B**. Collagen IV (A) and Vimentin (B) immunostaining of representative lung sections of 4-, 8-, 12-month-old *Sftpc^−/−^* mice and control mice. Higher magnification in the boxed area (n=3/group).

**Supplementary Figure 3:**
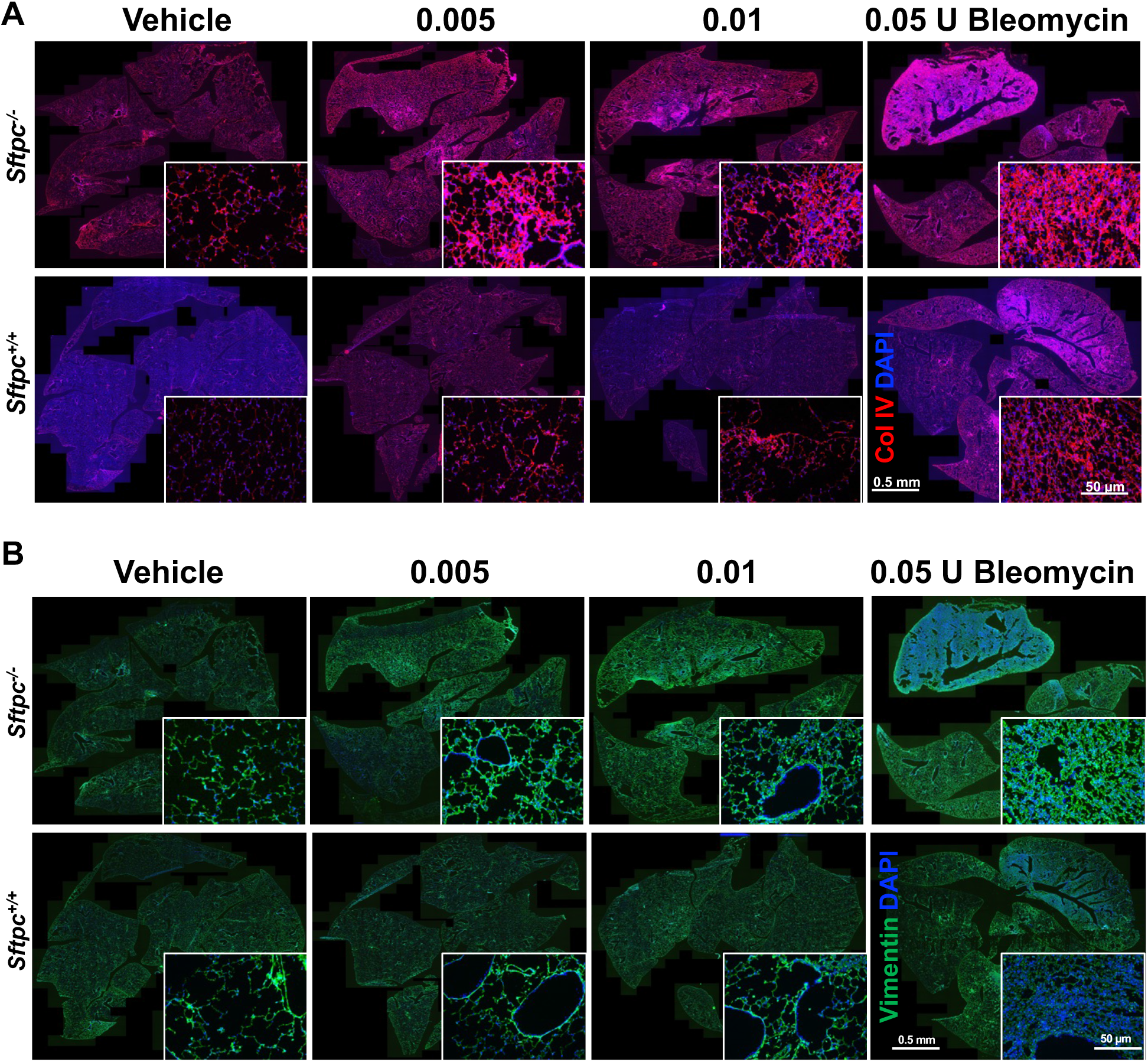
Analysis of lung parenchymal alteration in *Sftpc^−/−^* mice post bleomycin treatment. **A** and **B**. Collagen IV (A) and vimentin (B) immunostaining of representative lung sections of 4-month-old *Sftpc^−/−^* mice and control mice treated with bleomycin (n=3/group). Higher magnification represented in the boxed area.

**Supplementary Figure 4:**
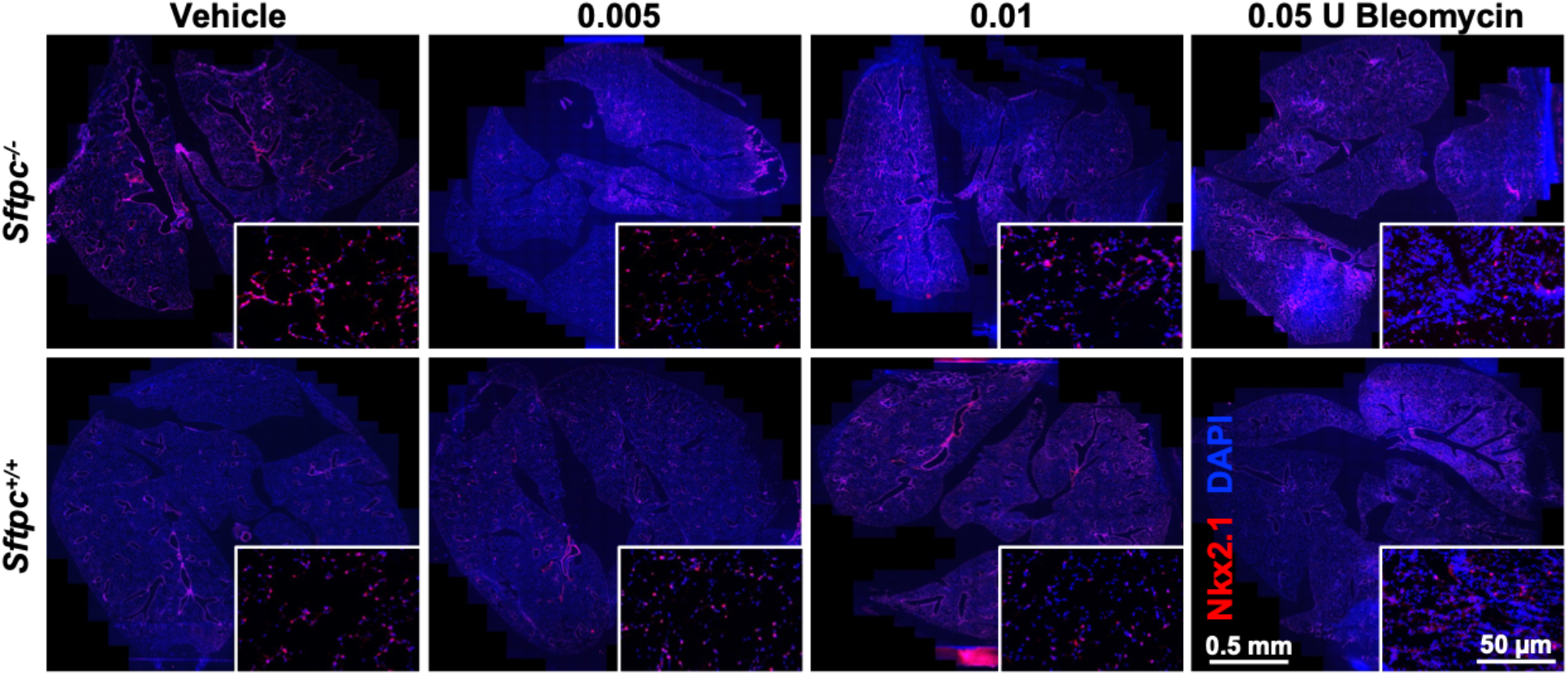
ATII cells immunostaining in *Sftpc*^−/−^ mice post bleomycin treatment. NKx2. 1 immunostaining of representative lung sections of 4-month-old *Sftpc^−/−^* and *Sftpc^−/−^* mice treated with increasing dose of bleomycin (0.005-0.05 U/mouse) (n=3.group). Higher magnification represented in the boxed area.

**Supplementary Figure 5:**
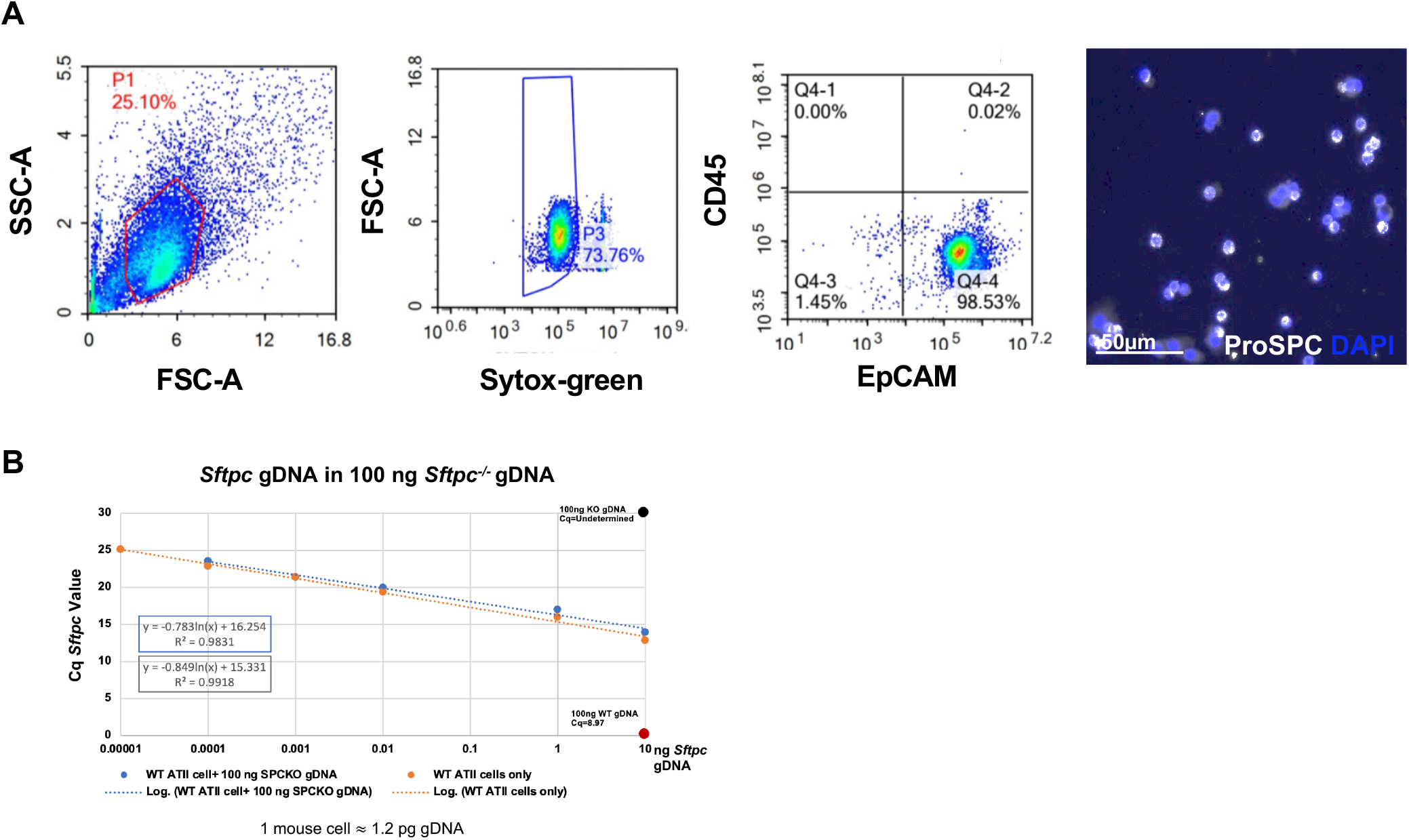
Isolation and purification of *Sftpc^+/+^* primary ATII cells. **(A)** Representative flow cytometric analysis for identification of ATII cells isolated after lung digestion and purification as described in Methods. ATII cells were identified by forward and side scatter followed by doublet discrimination of CD45 and EpCAM markers (n=7). On the right side, representative immunofluorescence staining for pro-SPC of ATII cells isolated as detailed in Methods and cultured overnight on Matrigel (1:30) to promote adherence. **(B)** Standard curve generated using the methodology for gDNA quantification previously described (56). Genomic DNA (nanograms) was extracted from increasing numbers of *Sftpc^+/+^* ATII cells sorted via FACS or lung tissue. DNA concentration and quality were assessed by absorbance (Nanodrop; Thermo Scientific, Wilmington, DE, USA). *Sftpc^+/+^* ATII cells gDNA samples alone or mixed with 100 ng *Sftpc^−/−^* lung tissue were subject to RT-qPCR for *Sftpc gene* to generate a standard curve. Cq values of 100 ng of gDNA isolated from *Sftpc^−/−^* and *Sftpc^+/+^* lung tissue were used as negative and positive controls, respectively. The line generated from this curve was used to calculate cell number values for all gDNA RT-qPCR experiments.

**Supplementary Figure 6:**
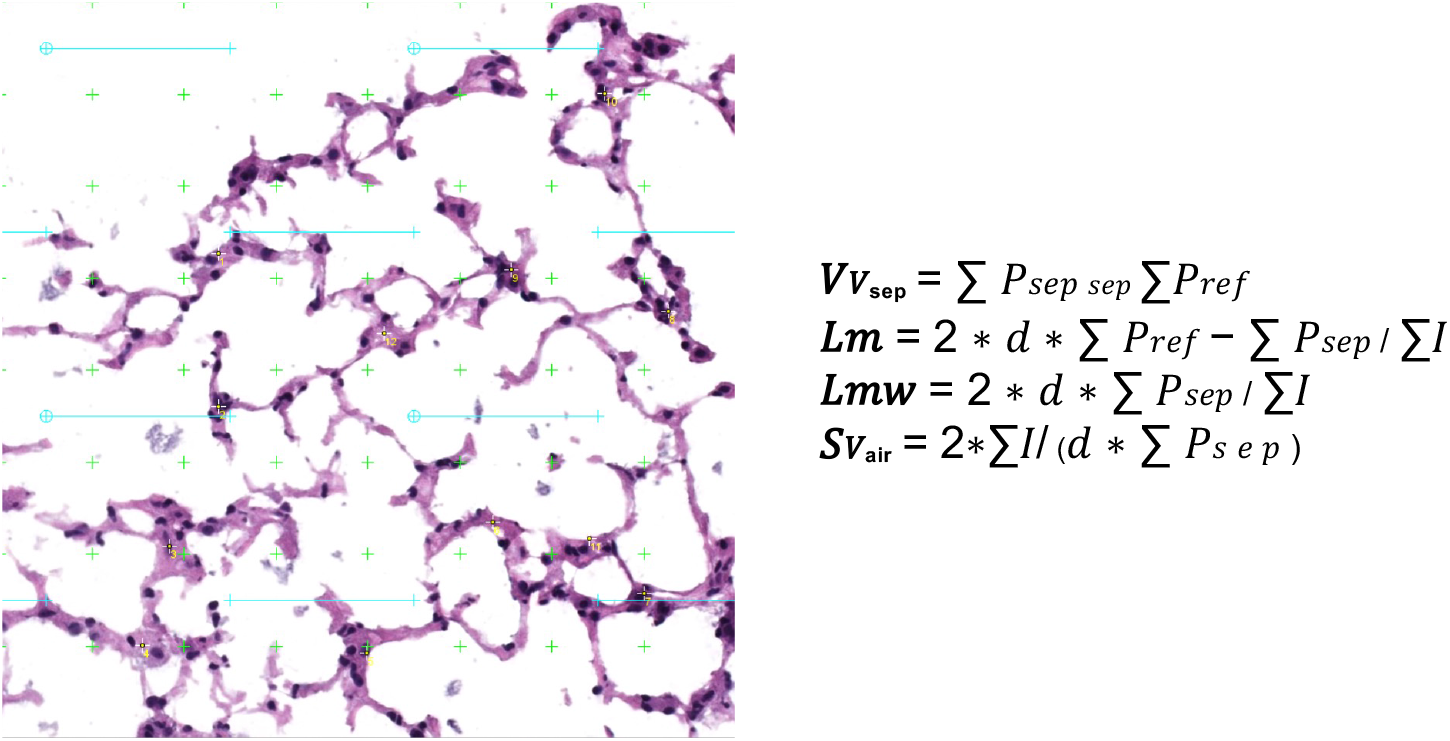
Representative grid and equations used in stereology analysis. A grid was randomly applied to 1000×1000 μm sections of lung histology and septal points were counted. Additionally, these same sections were analyzed using a semi-automated system and values were plugged into the given equations to determine these morphometric parameters.

**Supplementary Table 1:** Lung Injury Score. Lung Injury Score Criteria for scoring and calculation of lung injury score adapted from ATS guidelines (50,51).

Lung Injury Score
| Parameter | Score per Field |  |  |
| --- | --- | --- | --- |
|  | 0 | 1 | 2 |
| Neutrophils in the alveolar space | none | 1-5 | >5 |
| Neutrophils in the interstitial space | none | 1-5 | >5 |
| Hyaline membranes* | none | 1 occurrence | >1 distinct occurrence |
| Proteinaceous debris* | none | 1 occurrence | >1 distinct occurrence |
| Septal thickening | <2x | 2x-4x | >4x |

Total score per field = [(20 X A) + (14 X B) + (7 X C) + (7 X D) + (2 X E)]
\*In small animal models, hyaline membranes may be difficult to separate from proteinaceous debris.

**Supplementary Table 2.** Primary antibodies with dilutions. All antibodies were diluted in 5% Donkey Serum in PBS.

| Primary Antibody | Host Species | Company | Catalog Number | Perm. | Dilution |
| --- | --- | --- | --- | --- | --- |
| Anti-Pro-SPC | Rabbit | Seven Hills | WRAB9337 | Y | 1:400 |
| Anti-LAMP3 | Guinea Pig | Synaptic Signalling | 391005 | Y | 1:400 |
| Anti-Mature SPC | Rabbit | Seven Hills | WRAB76694 | Y | 1:400 |
| Anti-AGER/RAGE | Goat | R&D Systems | AF1145 | N | 1:200 |
| Anti-NKX2.1 | Rabbit | Seven Hills | WRAB1231 | Y | 1:500 |
| Anti-Collagen I | Rabbit | Abcam | ab34710 | N | 1:100 |
| Anti-Collagen IV | Rabbit | Abcam | Ab6586 | N | 1:500 |
| Anti-Vimentin | Rabbit | Abcam | ab92547 | Y | 1:400 |

**Supplementary Table 3.** Secondary Antibodies. Dilution for all secondary antibodies was 1:400 in 5% Donkey Serum in PBS.

| Secondary antibody | Host | Company | Catalog Number | Excitation Wavelength (Alexa Fluor) |
| --- | --- | --- | --- | --- |
| Anti-Goat IgG | Donkey | Invitrogen | A11057 | 568 |
| Anti-Rabbit IgG | Donkey | Invitrogen | A21206 | 488 |
| Anti-Rabbit IgG | Donkey | Invitrogen | A31572 | 555 |
| Anti-Rabbit IgG | Donkey | Invitrogen | A31573 | 647 |
| Anti-Guinea Pig IgG | Goat | Invitrogen | A21435 | 555 |

**Supplementary Table 4.**
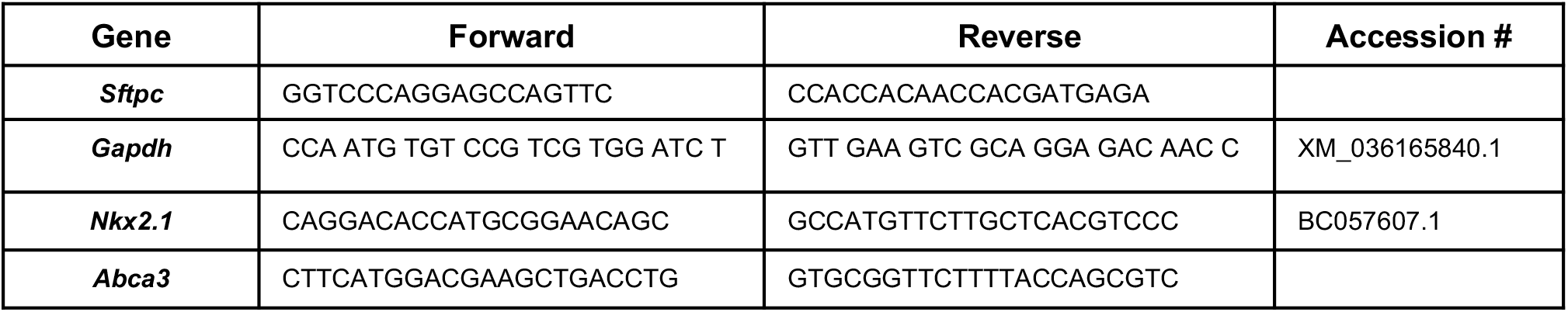
Primer sequences used for qPCR analysis.

## References

1. Gower WA, Nogee LM. Surfactant Dysfunction. Paediatric Respiratory Reviews. 2011;12(4):223–229. doi:10.1016/j.prrv.2011.01.005

2. Olmeda B, Martínez-Calle M, Pérez-Gil J. Pulmonary surfactant metabolism in the alveolar airspace: Biogenesis, extracellular conversions, recycling. Annals of Anatomy - Anatomischer Anzeiger. 2017;209:78–92. doi:10.1016/j.aanat.2016.09.008

3. Agassandian M, Mallampalli RK. Surfactant phospholipid metabolism. Biochimica et Biophysica Acta (BBA) - Molecular and Cell Biology of Lipids. 2013;1831(3):612–625. doi:10.1016/j.bbalip.2012.09.010

4. Parra E, Pérez-Gil J. Composition, structure and mechanical properties define performance of pulmonary surfactant membranes and films. Chemistry and Physics of Lipids. 2015;185:153–175. doi:10.1016/j.chemphyslip.2014.09.002

5. Rider ED, Ikegami M, Jobe AH. Localization of alveolar surfactant clearance in rabbit lung cells. American Journal of Physiology-Lung Cellular and Molecular Physiology. 1992;263(2):L201–L209. doi:10.1152/ajplung.1992.263.2.L201

6. Cunningham S, Jaffe A, Young LR. Children’s interstitial and diffuse lung disease. The Lancet Child & Adolescent Health. 2019;3(8):568–577. doi:10.1016/S2352-4642(19)30117-8

7. Singh J, Jaffe A, Schultz A, Selvadurai H. Surfactant protein disorders in childhood interstitial lung disease. Eur J Pediatr. 2021;180(9):2711–2721. doi:10.1007/s00431-021-04066-3

8. Fan LL, Dishop MK, Galambos C, et al. Diffuse Lung Disease in Biopsied Children 2 to 18 Years of Age. Application of the chILD Classification Scheme. Annals ATS. 2015;12(10):1498–1505. doi:10.1513/AnnalsATS.201501-064OC

9. Nogee LM. Genetic causes of surfactant protein abnormalities. Curr Opin Pediatr. 2019;31(3):330–339. doi:10.1097/MOP.0000000000000751

10. Kurland G, Deterding RR, Hagood JS, et al. An Official American Thoracic Society Clinical Practice Guideline: Classification, Evaluation, and Management of Childhood Interstitial Lung Disease in Infancy. Am J Respir Crit Care Med. 2013;188(3):376–394. doi:10.1164/rccm.201305-0923ST

11. Wang JY, Young LR. Insights into the Pathogenesis of Pulmonary Fibrosis from Genetic Diseases. Am J Respir Cell Mol Biol. 2022;67(1):20–35. doi:10.1165/rcmb.2021-0557TR

12. Griese M, Haug M, Brasch F, et al. Incidence and classification of pediatric diffuse parenchymal lung diseases in Germany. Orphanet J Rare Dis. 2009;4(1):26. doi:10.1186/1750-1172-4-26

13. Kornum JB, Christensen S, Grijota M, et al. The incidence of interstitial lung disease 1995– 2005: a Danish nationwide population-based study. BMC Pulm Med. 2008;8(1):24. doi:10.1186/1471-2466-8-24

14. Saddi V, Beggs S, Bennetts B, et al. Childhood interstitial lung diseases in immunocompetent children in Australia and New Zealand: a decade’s experience. Orphanet J Rare Dis. 2017;12(1):133. doi:10.1186/s13023-017-0637-x

15. Deutsch GH, Young LR, Deterding RR, et al. Diffuse lung disease in young children: application of a novel classification scheme. Am J Respir Crit Care Med. 2007;176(11):1120–1128. doi:10.1164/rccm.200703-393OC

16. Nathan N, Taam R, Epaud R, et al. A national internet-linked based database for pediatric interstitial lung diseases: the French network. Orphanet J Rare Dis. 2012;7(1):40. doi:10.1186/1750-1172-7-40

17. Nureki SI, Tomer Y, Venosa A, et al. Expression of mutant Sftpc in murine alveolar epithelia drives spontaneous lung fibrosis. Journal of Clinical Investigation. 2018;128(9):4008–4024. doi:10.1172/JCI99287

18. Katzen J, Beers MF. Contributions of alveolar epithelial cell quality control to pulmonary fibrosis. Journal of Clinical Investigation. 2020;130(10):5088–5099. doi:10.1172/JCI139519

19. Mulugeta S, Nureki SI, Beers MF. Lost after translation: insights from pulmonary surfactant for understanding the role of alveolar epithelial dysfunction and cellular quality control in fibrotic lung disease. Am J Physiol Lung Cell Mol Physiol. 2015;309(6):L507–525. doi:10.1152/ajplung.00139.2015

20. Beers MF, Morrisey EE. The three R’s of lung health and disease: repair, remodeling, and regeneration. J Clin Invest. 2011;121(6):2065–2073. doi:10.1172/JCI45961

21. Beers MF, Moodley Y. When Is an Alveolar Type 2 Cell an Alveolar Type 2 Cell? A Conundrum for Lung Stem Cell Biology and Regenerative Medicine. Am J Respir Cell Mol Biol. 2017;57(1):18–27. doi:10.1165/rcmb.2016-0426PS

22. Basil MC, Katzen J, Engler AE, et al. The Cellular and Physiological Basis for Lung Repair and Regeneration: Past, Present, and Future. Cell Stem Cell. 2020;26(4):482–502. doi:10.1016/j.stem.2020.03.009

23. Ruaro B, Salton F, Braga L, et al. The History and Mystery of Alveolar Epithelial Type II Cells: Focus on Their Physiologic and Pathologic Role in Lung. IJMS. 2021;22(5):2566. doi:10.3390/ijms22052566

24. Hogan BLM, Barkauskas CE, Chapman HA, et al. Repair and Regeneration of the Respiratory System: Complexity, Plasticity, and Mechanisms of Lung Stem Cell Function. Cell Stem Cell. 2014;15(2):123–138. doi:10.1016/j.stem.2014.07.012

25. Sisson TH, Mendez M, Choi K, et al. Targeted injury of type II alveolar epithelial cells induces pulmonary fibrosis. Am J Respir Crit Care Med. 2010;181(3):254–263. doi:10.1164/rccm.200810-1615OC

26. Garcia O, Hiatt MJ, Lundin A, et al. Targeted Type 2 Alveolar Cell Depletion. A Dynamic Functional Model for Lung Injury Repair. Am J Respir Cell Mol Biol. 2016;54(3):319–330. doi:10.1165/rcmb.2014-0246OC

27. Barkauskas CE, Cronce MJ, Rackley CR, et al. Type 2 alveolar cells are stem cells in adult lung. J Clin Invest. 2013;123(7):3025–3036. doi:10.1172/JCI68782

28. Deterding RR, DeBoer EM, Cidon MJ, et al. Approaching Clinical Trials in Childhood Interstitial Lung Disease and Pediatric Pulmonary Fibrosis. Am J Respir Crit Care Med. 2019;200(10):1219–1227. doi:10.1164/rccm.201903-0544CI

29. Benoit TM, Benden C. Pediatric lung transplantation: supply and demand. Curr Opin Organ Transplant. 2019;24(3):324–328. doi:10.1097/MOT.0000000000000630

30. Valapour M, Lehr CJ, Skeans MA, et al. OPTN/SRTR 2020 Annual Data Report: Lung. American J Transplantation. 2022;22(S2):438–518. doi:10.1111/ajt.16991

31. Alapati D, Zacharias WJ, Hartman HA, et al. In utero gene editing for monogenic lung disease. Sci Transl Med. 2019;11(488):eaav8375. doi:10.1126/scitranslmed.aav8375

32. Serrano-Mollar A, Nacher M, Gay-Jordi G, Closa D, Xaubet A, Bulbena O. Intratracheal transplantation of alveolar type II cells reverses bleomycin-induced lung fibrosis. Am J Respir Crit Care Med. 2007;176(12):1261–1268. doi:10.1164/rccm.200610-1491OC

33. Guillamat-Prats R, Gay-Jordi G, Xaubet A, Peinado VI, Serrano-Mollar A. Alveolar Type II cell transplantation restores pulmonary surfactant protein levels in lung fibrosis. The Journal of Heart and Lung Transplantation. 2014;33(7):758–765. doi:10.1016/j.healun.2014.03.008

34. Weiner AI, Jackson SR, Zhao G, et al. Mesenchyme-free expansion and transplantation of adult alveolar progenitor cells: steps toward cell-based regenerative therapies. npj Regen Med. 2019;4(1):17. doi:10.1038/s41536-019-0080-9

35. Walters MC, Patience M, Leisenring W, et al. Stable mixed hematopoietic chimerism after bone marrow transplantation for sickle cell anemia. Biology of Blood and Marrow Transplantation. 2001;7(12):665–673. doi:10.1053/bbmt.2001.v7.pm11787529

36. Fitzhugh CD, Cordes S, Taylor T, et al. At least 20% donor myeloid chimerism is necessary to reverse the sickle phenotype after allogeneic HSCT. Blood. 2017;130(17):1946–1948. doi:10.1182/blood-2017-03-772392

37. Okaygoun D, Oliveira DD, Soman S, Williams R. Advances in the management of haemophilia: emerging treatments and their mechanisms. J Biomed Sci. 2021;28(1):64. doi:10.1186/s12929-021-00760-4

38. Meena NK, Ralston E, Raben N, Puertollano R. Enzyme Replacement Therapy Can Reverse Pathogenic Cascade in Pompe Disease. Molecular Therapy - Methods & Clinical Development. 2020;18:199–214. doi:10.1016/j.omtm.2020.05.026

39. Whitsett JA, Weaver TE. Hydrophobic Surfactant Proteins in Lung Function and Disease. N Engl J Med. 2002;347(26):2141–2148. doi:10.1056/NEJMra022387

40. Nogee LM, Dunbar AE, Wert S, Askin F, Hamvas A, Whitsett JA. Mutations in the Surfactant Protein C Gene Associated With Interstitial Lung Disease. Chest. 2002;121(3):20S–21S. doi:10.1378/chest.121.3_suppl.20S

41. Brasch F, Griese M, Tredano M, et al. Interstitial lung disease in a baby with a de novo mutation in the SFTPC gene. European Respiratory Journal. 2004;24(1):30–39. doi:10.1183/09031936.04.00000104

42. Wambach JA, Yang P, Wegner DJ, et al. Surfactant Protein-C Promoter Variants Associated With Neonatal Respiratory Distress Syndrome Reduce Transcription. Pediatr Res. 2010;68(3):216–220. doi:10.1203/PDR.0b013e3181eb5d68

43. Amin RS, Wert SE, Baughman RP, et al. Surfactant protein deficiency in familial interstitial lung disease. The Journal of Pediatrics. 2001;139(1):85–92. doi:10.1067/mpd.2001.114545

44. Tredano M, Griese M, Brasch F, et al. Mutation of *SFTPC* in infantile pulmonary alveolar proteinosis with or without fibrosing lung disease. Am J Med Genet. 2004;126A(1):18–26. doi:10.1002/ajmg.a.20670

45. Liptzin DR, Patel T, Deterding RR. Chronic Ventilation in Infants with Surfactant Protein C Mutations: An Alternative to Lung Transplantation. Am J Respir Crit Care Med. 2015;191(11):1338–1340. doi:10.1164/rccm.201411-1955LE

46. Thomas AQ, Lane K, Phillips J, et al. Heterozygosity for a Surfactant Protein C Gene Mutation Associated with Usual Interstitial Pneumonitis and Cellular Nonspecific Interstitial Pneumonitis in One Kindred. Am J Respir Crit Care Med. 2002;165(9):1322–1328. doi:10.1164/rccm.200112-123OC

47. Glasser SW, Detmer EA, Ikegami M, Na CL, Stahlman MT, Whitsett JA. Pneumonitis and emphysema in sp-C gene targeted mice. J Biol Chem. 2003;278(16):14291–14298. doi:10.1074/jbc.M210909200

48. Toth A, Steinmeyer S, Kannan P, et al. Inflammatory blockade prevents injury to the developing pulmonary gas exchange surface in preterm primates. Sci Transl Med. 2022;14(638):eabl8574. doi:10.1126/scitranslmed.abl8574

49. Dorrello NV, Guenthart BA, O’Neill JD, et al. Functional vascularized lung grafts for lung bioengineering. Sci Adv. 2017;3(8):e1700521. doi:10.1126/sciadv.1700521

50. Kulkarni HS, Lee JS, Bastarache JA, et al. Update on the Features and Measurements of Experimental Acute Lung Injury in Animals: An Official American Thoracic Society Workshop Report. Am J Respir Cell Mol Biol. 2022;66(2):e1–e14. doi:10.1165/rcmb.2021-0531ST

51. Matute-Bello G, Downey G, Moore BB, et al. An Official American Thoracic Society Workshop Report: Features and Measurements of Experimental Acute Lung Injury in Animals. Am J Respir Cell Mol Biol. 2011;44(5):725–738. doi:10.1165/rcmb.2009-0210ST

52. Ruwisch J, Sehlmeyer K, Roldan N, et al. Air Space Distension Precedes Spontaneous Fibrotic Remodeling and Impaired Cholesterol Metabolism in the Absence of Surfactant Protein C. Am J Respir Cell Mol Biol. 2020;62(4):466–478. doi:10.1165/rcmb.2019-0358OC

53. Glasser SW, Burhans MS, Korfhagen TR, et al. Altered stability of pulmonary surfactant in SP-C-deficient mice. Proceedings of the National Academy of Sciences. 2001;98(11):6366–6371. doi:10.1073/pnas.101500298

54. Lawson WE, Polosukhin VV, Stathopoulos GT, et al. Increased and prolonged pulmonary fibrosis in surfactant protein C-deficient mice following intratracheal bleomycin. Am J Pathol. 2005;167(5):1267–1277. doi:10.1016/S0002-9440(10)61214-X

55. Madala SK, Maxfield MD, Davidson CR, et al. Rapamycin Regulates Bleomycin-Induced Lung Damage in SP-C-Deficient Mice. Pulmonary Medicine. 2011;2011:1–12. doi:10.1155/2011/653524

56. Prigent J, Herrero A, Ambroise J, Smets F, Deblandre GA, Sokal EM. Human Progenitor Cell Quantification after Xenotransplantation in Rat and Mouse Models by a Sensitive qPCR Assay. Cell Transplant. 2015;24(8):1639–1652. doi:10.3727/096368914X681955

57. Glasser SW, Senft AP, Whitsett JA, et al. Macrophage Dysfunction and Susceptibility to Pulmonary *Pseudomonas aeruginosa* Infection in Surfactant Protein C-Deficient Mice. J Immunol. 2008;181(1):621–628. doi:10.4049/jimmunol.181.1.621

58. Glasser SW, Witt TL, Senft AP, et al. Surfactant protein C-deficient mice are susceptible to respiratory syncytial virus infection. American Journal of Physiology-Lung Cellular and Molecular Physiology. 2009;297(1):L64–L72. doi:10.1152/ajplung.90640.2008

59. Glasser SW, Maxfield MD, Ruetschilling TL, et al. Persistence of LPS-Induced Lung Inflammation in Surfactant Protein-C–Deficient Mice. Am J Respir Cell Mol Biol. 2013;49(5):845–854. doi:10.1165/rcmb.2012-0374OC

60. Venosa A, Katzen J, Tomer Y, et al. Epithelial Expression of an Interstitial Lung Disease– Associated Mutation in Surfactant Protein-C Modulates Recruitment and Activation of Key Myeloid Cell Populations in Mice. JI. 2019;202(9):2760–2771. doi:10.4049/jimmunol.1900039

61. Nogee LM. Genetic causes of surfactant protein abnormalities: Current Opinion in Pediatrics. 2019;31(3):330–339. doi:10.1097/MOP.0000000000000751

62. Cameron HS, Somaschini M, Carrera P, et al. A common mutation in the surfactant protein C gene associated with lung disease. The Journal of Pediatrics. 2005;146(3):370–375. doi:10.1016/j.jpeds.2004.10.028

63. Sehlmeyer K, Ruwisch J, Roldan N, Lopez-Rodriguez E. Alveolar Dynamics and Beyond – The Importance of Surfactant Protein C and Cholesterol in Lung Homeostasis and Fibrosis. Front Physiol. 2020;11:386. doi:10.3389/fphys.2020.00386

64. Katzen J, Wagner BD, Venosa A, et al. An SFTPC BRICHOS mutant links epithelial ER stress and spontaneous lung fibrosis. JCI Insight. 2019;4(6). doi:10.1172/jci.insight.126125

65. Bridges JP, Wert SE, Nogee LM, Weaver TE. Expression of a Human Surfactant Protein C Mutation Associated with Interstitial Lung Disease Disrupts Lung Development in Transgenic Mice. Journal of Biological Chemistry. 2003;278(52):52739–52746. doi:10.1074/jbc.M309599200

66. Lawson WE, Cheng DS, Degryse AL, et al. Endoplasmic reticulum stress enhances fibrotic remodeling in the lungs. Proc Natl Acad Sci USA. 2011;108(26):10562–10567. doi:10.1073/pnas.1107559108

67. Sitaraman S, Martin EP, Na CL, et al. Surfactant protein C mutation links postnatal type 2 cell dysfunction to adult disease. JCI Insight. 2021;6(14):e142501. doi:10.1172/jci.insight.142501

68. O’Dwyer DN, Moore BB. Animal Models of Pulmonary Fibrosis. In: Alper S, Janssen WJ, eds. Lung Innate Immunity and Inflammation. Vol 1809. Methods in Molecular Biology. Springer New York; 2018:363–378. doi:10.1007/978-1-4939-8570-8_24

69. Milman Krentsis I, Rosen C, Shezen E, et al. Lung Injury Repair by Transplantation of Adult Lung Cells Following Preconditioning of Recipient Mice. Stem Cells Translational Medicine. 2018;7(1):68–77. doi:10.1002/sctm.17-0149

70. Rosen C, Shezen E, Aronovich A, et al. Preconditioning allows engraftment of mouse and human embryonic lung cells, enabling lung repair in mice. Nat Med. 2015;21(8):869–879. doi:10.1038/nm.3889

71. Herriges MJ, Yampolskaya M, Thapa BR, et al. Durable Alveolar Engraftment of PSC-Derived Lung Epithelial Cells into Immunocompetent Mice. Cell Biology; 2022. doi:10.1101/2022.07.26.501591

72. Kathiriya JJ, Brumwell AN, Jackson JR, Tang X, Chapman HA. Distinct Airway Epithelial Stem Cells Hide among Club Cells but Mobilize to Promote Alveolar Regeneration. Cell Stem Cell. 2020;26(3):346–358.e4. doi:10.1016/j.stem.2019.12.014

73. Louie SM, Moye AL, Wong IG, et al. Progenitor potential of lung epithelial organoid cells in a transplantation model. Cell Reports. 2022;39(2):110662. doi:10.1016/j.celrep.2022.110662

74. Vaughan AE, Brumwell AN, Xi Y, et al. Lineage-negative progenitors mobilize to regenerate lung epithelium after major injury. Nature. 2015;517(7536):621–625. doi:10.1038/nature14112

75. Miller AJ, Hill DR, Nagy MS, et al. In Vitro Induction and In Vivo Engraftment of Lung Bud Tip Progenitor Cells Derived from Human Pluripotent Stem Cells. Stem Cell Reports. 2018;10(1):101–119. doi:10.1016/j.stemcr.2017.11.012

76. Nichane M, Javed A, Sivakamasundari V, et al. Isolation and 3D expansion of multipotent Sox9+ mouse lung progenitors. Nat Methods. 2017;14(12):1205–1212. doi:10.1038/nmeth.4498

77. Melton KR, Nesslein LL, Ikegami M, et al. SP-B deficiency causes respiratory failure in adult mice. American Journal of Physiology-Lung Cellular and Molecular Physiology. 2003;285(3):L543–L549. doi:10.1152/ajplung.00011.2003

78. Dzhuraev G, Rodríguez-Castillo JA, Ruiz-Camp J, et al. Estimation of absolute number of alveolar epithelial type 2 cells in mouse lungs: a comparison between stereology and flow cytometry. Journal of Microscopy. 2019;275(1):36–50. doi:10.1111/jmi.12800

79. Lederer, David J, Martinez, Fernando J. Idiopathic Pulmonary Fibrosis. N Engl J Med. 2018;379(8):795–798. doi:10.1056/NEJMc1807508

80. Chen YW, Huang SX, de Carvalho ALRT, et al. A three-dimensional model of human lung development and disease from pluripotent stem cells. Nat Cell Biol. 2017;19(5):542–549. doi:10.1038/ncb3510

81. Jacob A, Vedaie M, Roberts DA, et al. Derivation of self-renewing lung alveolar epithelial type II cells from human pluripotent stem cells. Nat Protoc. 2019;14(12):3303–3332. doi:10.1038/s41596-019-0220-0

82. Zacharias WJ, Frank DB, Zepp JA, et al. Regeneration of the lung alveolus by an evolutionarily conserved epithelial progenitor. Nature. 2018;555(7695):251–255. doi:10.1038/nature25786

83. Salaets T, Tack B, Gie A, et al. A semi-automated method for unbiased alveolar morphometry: Validation in a bronchopulmonary dysplasia model. Koval M, ed. PLoS ONE. 2020;15(9):e0239562. doi:10.1371/journal.pone.0239562

